# Multiomic Immunophenotyping of COVID-19 Patients Reveals Early Infection Trajectories

**DOI:** 10.1101/2020.07.27.224063

**Authors:** Yapeng Su, Daniel Chen, Christopher Lausted, Dan Yuan, Jongchan Choi, Cheng Dai, Valentin Voillet, Kelsey Scherler, Pamela Troisch, Venkata R. Duvvuri, Priyanka Baloni, Guangrong Qin, Brett Smith, Sergey Kornilov, Clifford Rostomily, Alex Xu, Jing Li, Shen Dong, Alissa Rothchild, Jing Zhou, Kim Murray, Rick Edmark, Sunga Hong, Lesley Jones, Yong Zhou, Ryan Roper, Sean Mackay, D. Shane O’Mahony, Christopher R Dale, Julie A Wallick, Heather A Algren, Zager A Michael, Andrew Magis, Wei Wei, Nathan D. Price, Sui Huang, Naeha Subramanian, Kai Wang, Jennifer Hadlock, Leroy Hood, Alan Aderem, Jeffrey A. Bluestone, Lewis L. Lanier, Phil Greenberg, Raphael Gottardo, Mark M. Davis, Jason D. Goldman, James R. Heath, the ISB-Swedish COVID19 Biobanking Unit

## Abstract

Host immune responses play central roles in controlling SARS-CoV2 infection, yet remain incompletely characterized and understood. Here, we present a comprehensive immune response map spanning 454 proteins and 847 metabolites in plasma integrated with single-cell multi-omic assays of PBMCs in which whole transcriptome, 192 surface proteins, and T and B cell receptor sequence were co-analyzed within the context of clinical measures from 50 COVID19 patient samples. Our study reveals novel cellular subpopulations, such as proliferative exhausted CD8^+^ and CD4^+^ T cells, and cytotoxic CD4^+^ T cells, that may be features of severe COVID-19 infection. We condensed over 1 million immune features into a single immune response axis that independently aligns with many clinical features and is also strongly associated with disease severity. Our study represents an important resource towards understanding the heterogeneous immune responses of COVID-19 patients and may provide key information for informing therapeutic development.

## INTRODUCTION

The novel coronavirus disease, COVID-19 has rapidly spread to become a global health challenge, with over 11M cases and over 0.5M associated fatalities reported by early July 2020. 20-31% of symptomatic patients require hospitalization, with ICU admission rates ranging from 4.9-11.5%, and fatality rates ranging from 2-10% (Iype and Gulati, 2020). Management of COVID-19 is mostly supportive, and respiratory failure from acute respiratory distress syndrome (ARDS) is the leading cause of mortality. Patients with more severe COVID-19 infections are distinguished by significant immune dysregulation, the nature of which is incompletely understood.

Recorded signatures of immune dysregulation include lymphopenia that is commonly associated with reduced numbers of T cells, natural killer cells, and other lymphocytes (Cao, 2020; Ruan et al., 2020; Tan et al., 2020), as well as elevated exhaustion markers on immune cells (Zheng et al., 2020). Elevated levels of circulating inflammatory cytokines (Mathew et al., 2020) have been reported, although the primary source of such cytokines, as well as the roles of circulating T cells and monocytes, has been debated (Wilk et al., 2020; Zhou et al., 2020). A condition bearing similarities to IL-6-mediated cytokine release syndrome (CRS) (Yang et al., 2020) has also been reported in severely ill COVID-19 patients, although unlike in CRS in cancer patients receiving CAR-T immunotherapies, associated toxicity are less severe, and benefit of IL-6 blockade was less consistent (Vardhana and Wolchok, 2020). However, high IL-6 levels, together with elevated levels of C-reactive protein, has been shown to correlate with the need for mechanical ventilation (Herold et al., 2020; Wang et al., 2020).

Single-cell molecular profiling has just begun to unravel the heterogeneous nature of immune dysregulation in COVID-19 patients (Herold et al., 2020; Wang et al., 2020). These have included reports that airway epithelium-immune cell interactions increase with increasing disease severity, potentially providing insights into lung injury and the respiratory failure (Chua et al., 2020). Wilk et al. reported on the sc-RNA-seq profiles of PBMCs collected from seven COVID-19 patients (Wilk et al., 2020) to yield a preliminary single cell PBMC analysis, which revealed factors such as a heterogeneous interferon-stimulated gene signature and robust HLA Class II downregulation. This study also revealed a high patient-to-patient variability, emphasizing the need for more comprehensive studies. Mathew et al. reported an analysis of ~200 immune cell features, integrated with >30 clinical features, for 71 COVID-19 patients (Mathew et al., 2020) to resolve immune cell phenotypes associated with both poor and improving clinical status in COVID-19 patients. This larger sample size allowed for the identification of certain canonical immune cell phenotypes that are associated with COVID-19 infection and clinical features of the disease. These findings collectively suggest that approaches that can connect information levels ranging from clinical characteristics to immune features at single-cell resolution may well be necessary for accelerating our understanding of the broad clinical heterogeneity of SARS-CoV2 infections.

Here we develop a comprehensive view of the 1-week period of COVID-19 infection following initial hospitalization, for a cohort of 26 hospitalized patients compared with 35 healthy donors. This view is built by first extracting the demographics and static and dynamic clinical features for each patient from their electronic health record. This clinical picture is then integrated with multi-omic analytics of two sequential blood draws per patient (50 total), the first of which was collected shortly after initial clinical diagnosis (time = T1), and the second a few days later (time = T2) (**Figure S1A**). For each blood draw, the plasma levels of nearly 500 proteins and 1000 metabolites were quantified **(Figure 1A)**. That data set was complemented by single-cell multi-omic analyses of PBMCs in which the whole transcriptome, 192 surface proteins, 32 secreted proteins, and T-cell and B-cell receptor gene sequences are measured. Our results not only validated past studies on correlations between distinct subpopulation presence and disease severity but also revealed many novel COVID-19 associated immune cell subpopulations and functions, many of which associate with disease severity. Our comprehensive clinical, multi-omic single-cell and plasma analyte datasets provide an integrated multi-omic resource of the immune response to severe COVID-19 infection. Our unique resource can permit the elucidation of disease perturbed, coordinated biological networks that capture the disease process, which are not readily resolved using less comprehensive approaches.

**Figure 1.**
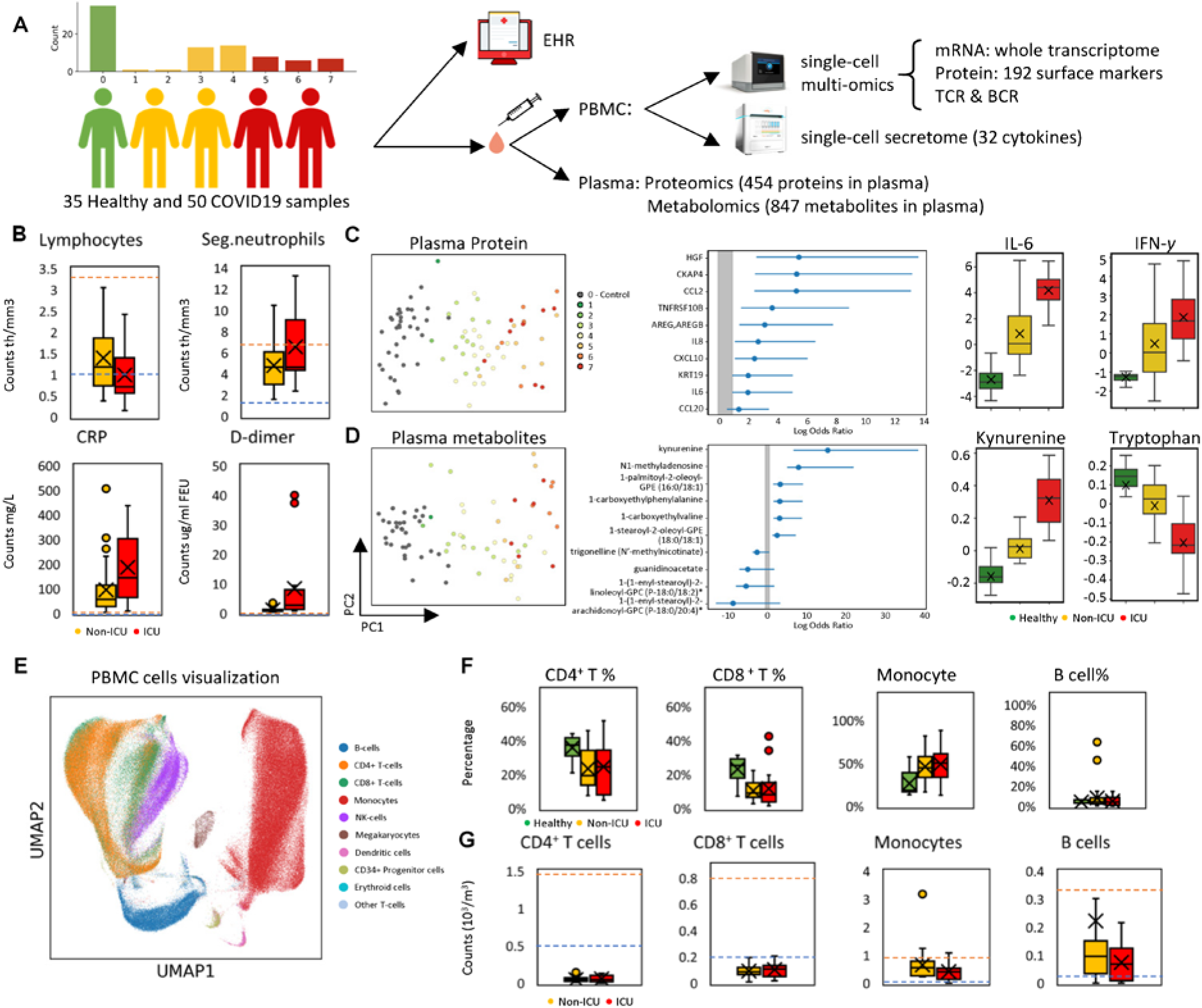
Overview of the multi-omic characterization of immune responses in COVID-19 patients. A. Overview of the Swedish/ISB INCOV study of COVID-19 patients. The bar graph represents the disease severity, using the WHO ordinal score, of (hospitalized) patients studied through detailed analyses of PBMCs and blood plasma, collected near the time of diagnosis (T1), and approximately 1 week later (T2). PBMCs were characterized using 10x genomics and Isoplexis single cell methods. The 10X analysis simultaneously profiled the whole transcriptome, 192 surface proteins, and T cell and B cell receptor sequences. Isoplexis analyzes for a 32-plex secretome at single cell resolution from selected immune cell phenotypes. Plasma was analyzes using O-link technology to quantify the levels of 454 proteins, and Metabolon technology to assay for the levels of 847 metabolites. These data were integrated with clinical data from electronic health record for detailed characterization of the patients. B. Levels of selected clinical parameters comparing non-ICU (yellow) and ICU (red) COVID-19 samples. Ranges from healthy donors are below the orange and above the blue-dashed lines. Sample numbers analyzed were non-ICU (n=34) and ICU (n=16). C. Plasma protein analysis. Left panel: PCA analysis of 454 plasma proteins measured from 85 blood draws. Each dot is a single patient/healthy sample, with a color that corresponds to disease severity (see key). Middle panel: Forest plots depicting the odds ratios obtained from logistic regression analysis between cytokines and WHO ordinal scale. Grey shading indicates plot areas with odds ratios <1, indicating the normal range for non-COVID healthy control (defined by likelihood). Odds ratios are indicated with points, and confidence lines encompass the range between the lower and upper limits. Right panel: Box plot of protein concentrations in healthy donors (green), non-ICU blood draws (yellow), and ICU blood draws (red) of COVID-19 patients. D. Analysis of plasma metabolite data (847 metabolites from 85 samples). Left panel: PCA analysis of all samples’ metabolite data. Each dot represent a single sample, with a color that corresponds to the WHO ordinal scale for disease severity. Middle panel: Forest plots depicting the odds ratios obtained from logistic regression analysis between cytokines and disease severity. Grey shading indicates plot areas with odds ratios <1, indicating the normal range for non-COVID healthy control (defined by likelihood). Odds ratios are indicated with points, and confidence lines encompass the range between the lower and upper limits. Right panel: Box plot of metabolite concentrations in healthy donors (green), non-ICU blood draws (yellow), and ICU blood draws (red) of COVID-19 patients. E. 2D projection of all PBMCs from all samples using UMAP (Uniform Manifold Approximation and Projection). Single cells are shown as dots, colored by their cell type assignments. F. Box plot depicting the percentages of major immune cell types among PBMCs from healthy donors (green), non-ICU blood draws (yellow), and ICU blood draws (red) of COVID-19 patients. G. Box plot depicting the absolute count of major immune cell types from non-ICU blood draws (yellow) and ICU blood draws (red) of COVID-19 patients. Ranges that specify normal limits are indicated by the blue (lower) and orange (upper) dashed lines.

## RESULTS

### COVID-19 patients display varying clinical profiles, plasma proteomic and metabolomic profiles as well as circulating immune cell populations

To comprehensively characterize the peripheral immune response in severe COVID-19, we analyzed peripheral blood mononuclear cells (PBMCs) and plasma from 35 healthy and 50 COVID-19 patient samples (about 5000 cells per sample) (**Figures 1A and S1A**) To capture the phenotypes and functional properties of PBMCs, we applied 10X Genomics’ droplet-based single-cell multi-omic technology (Zheng et al., 2017) to simultaneously measure whole transcriptome (over 25000 genes), the abundance of 192 surface proteins and T and B cell receptor (TCR and BCR) sequences from each single cell. Further, we utilized the Isoplexis assay (Lu et al., 2015; Ma et al., 2011) to assess the levels of 32 secreted chemokines and cytokines from viable, single CD4^+^, CD8^+^ T cells and monocytes. In addition, plasma derived from the same patient samples were characterized by quantifying the levels of 454 targeted proteins using the Olink proximity extension assays and 847 untargeted metabolites from Metabolon’s global metabolomics platform. **(Figure 1A)**

These COVID-19 patients exhibited a wide range of clinical laboratory phenotypes, including many that correlate with disease severity, as defined by the WHO ordinal scale (WHO) (WHO, 2020) **(Figures 1B, S1B and S1C)**. Clinical metadata for these COVID-19 patient samples are presented in **Table S1**. For example, segmented neutrophil (Seg. neutrophils) numbers displayed a positive correlation with disease severity **(Figures 1B and S1C)**, as gauged by samples collected from patients treated in the intensive care unit (ICU) vs. those not in the ICU (**Table S1**). Lymphocyte count was reduced in ICU samples **(Figure 1B)**, consistent with previous reports of lymphopenia in severely infected COVID-19 patients (Cao et al., 2020). Furthermore, clinical inflammatory syndrome related features, including C reactive protein (CRP), ferritin, D-dimer, and fibrinogen, were all elevated in COVID-19 patients **(Figures 1B, and S1B)**, with the increase of CRP and D-dimer even more pronounced in ICU samples **(Figure 1B)**. We identified a number of other clinical features that also correlate with disease severity **(Figure S1C)**, and are consistent with existing literature (Mathew et al., 2020).

The analyzed plasma proteins and metabolites are listed in **Tables S2 and S3**. Principal component analysis (PCA) of both datasets revealed a clear separation of healthy and COVID-19 samples, with principal component 1 (PC1) positively correlated with disease severity **(Figures 1C, 1D and S1D)**. We further quantified the correlation of each protein or metabolite with disease severity. Consistent with previous reports (Somers et al., 2020), several cytokines associated with immune defense and inflammation, including IFN-γ, IL6, and IL8 showed positive correlations with disease severity and exhibited the highest levels in ICU samples **(Table S4, Figures 1C and S1E)**. Similarly, the inflammation and hypoxia associated metabolite lactate (Ivashkiv, 2020) exhibited a positive correlation with disease severity (**Table S5**). In contrast, trigonelline, uridine, and guanidinoacetate all displayed negative correlation **(Table S5, Figure 1D)**, indicating reduced antioxidant activity, increased superoxide dismutase activity, and altered antiinflammatory activity in COVID-19 patients (Cicko et al., 2015; Liu et al., 2018; Marques et al., 2019; Zhou et al., 2013). Interestingly, the immune suppressive metabolite kynurenine (Belladonna et al., 2009) showed the highest level in ICU samples while its precursor, tryptophan, showed the lowest level **(Figure 1D)**, suggesting an accelerated conversion of tryptophan towards kynurenine in these patients. These results highlight the perturbation of COVID-19 on metabolic functions, particularly upregulated immunosuppressive pathways, which may reduce the capacity of immune system to clear the virus in those patients. These broad changes of plasma proteomic and metabolomic profiles suggest a shift in biological programs associated with disease severity in COVID-19 patients.

We comprehensively characterized all PBMC immune cells using single-cell multi-omic analysis that encompasses the whole transcriptome and 192 surface proteins (**Table S6**) from each single cell. The resulting high-dimensional single-cell dataset was visualized as a two-dimensional map by Uniform Manifold Approximation and Projection (UMAP) (Becht et al., 2019). Cellular phenotype identities were revealed by unsupervised clustering followed by annotation with canonical marker genes and proteins for each major immune cell type **(Figures 1E, S1F, S1G and Methods)**. We then quantified the cell type proportions and their correlation with disease severity and observed significant variability across samples **(Figures 1F, S1H and S1I)**. Consistent with previous reports (Mathew et al., 2020), a drop in the T cell fraction, especially CD8^+^ T cells, relative to healthy donors was observed **(Figure 1F**). This pattern was also seen in the reduced absolute count of CD4^+^ and CD8^+^ T cells **(Figure 1G)**. By contrast, monocyte, NK cell and megakaryocyte percentages were elevated **(Figures 1F and S1I)**. In summary, high-dimensional multiomic analysis revealed an increase of inflammatory cytokines and potentially immunosuppressive metabolites in plasma, as well as significant alterations in the composition of major immune cell classes, all of which associated with disease severity. This prompted us to further investigate the complete transcriptomes of all immune cell types and their correlation with disease.

### Severity of COVID-19 is associated with the activation, proliferation, and clonal expansion of CD8^+^ T cells

The decrease in the fraction of circulating CD8^+^ T cells with an increase of SARS-CoV-2 infection severity prompted us to further investigate this immune cell class by studying its subpopulation composition, clonal expansion, and polyfunctionality. We first projected all CD8^+^ T cells onto a two dimensional UMAP and utilized unsupervised clustering of the single-cell whole transcriptomes to resolve 10 subpopulations **(Figure 2A, top left panel)**. Each subpopulation could be further characterized at the protein level **(Figure 2B, 2C, and S2B)** and by gene signatures through transcriptome analysis **(Figure 2C)**. The UMAP analysis revealed a clear spatial separation between the major CD8^+^ T cell phenotypes **(Figures 2A–2C),** which was independently validated using measured protein levels. In the individual panels of Figure 2A, we have used the expression of specific transcripts to identify naïve, memory, effector, exhausted-like, and proliferative phenotypes. For instance, the naïve cell mRNA and protein markers *LEF1, TCF7* and CD197 were all upregulated in the orange-colored cluster 1, while activated effector markers such as *GZMB* and *PRF1* were much higher in clusters 0 and 2 **(Figures 2A, 2C and S2A)**. The markers *GZMK* and *CD69* for memory-like (or intermediate-like) cells were mainly upregulated in cluster 3 (red), which is sandwiched between the regions of naïve and effector cells **(Figures 2A and S2A)**. Interestingly, cluster 8 (light blue) displayed intermediate levels of effector markers, but upregulated proliferation markers *MKI67* and *TYMS* **(Figures 2A and S2A)**. Exhausted CD8^+^ T cells have been documented in COVID-19 patients (Diao et al., 2020; Moon, 2020; Zheng et al., 2020) and, indeed, exhaustion markers *LAG3* and *TIGIT* were upregulated in both effector clusters (0 and 2) as well as the proliferative cluster 8 **(Figures 2A, 2C, S2A and S2B)**. The measured protein levels yielded a consistent picture. For example, the naïve-like cluster 1 displayed a higher CD45RA/CD45RO ratio than other clusters **(Figure 2B)**. As might be expected, the effector-like clusters 0 and 2 also showed the most clonal expansion **(Figures 2D, 2E, S2C and S2D)**, while T cells with a clonal expansion index of 1 were mainly identified within the naïve-like cluster 1 and the intermediate memory-like cluster 3 **(Figures 2D, 2E, S2C and S2D)**. This suggests that the activated effector CD8^+^ T cells detected in peripheral blood are clonally expanding due to virus-antigen encounters. Thus, the peripheral CD8^+^ T cells are heterogeneous, comprised of diverse subpopulations with distinct gene expression, surface protein profiles, and clonal-expansion levels.

**Figure 2.**
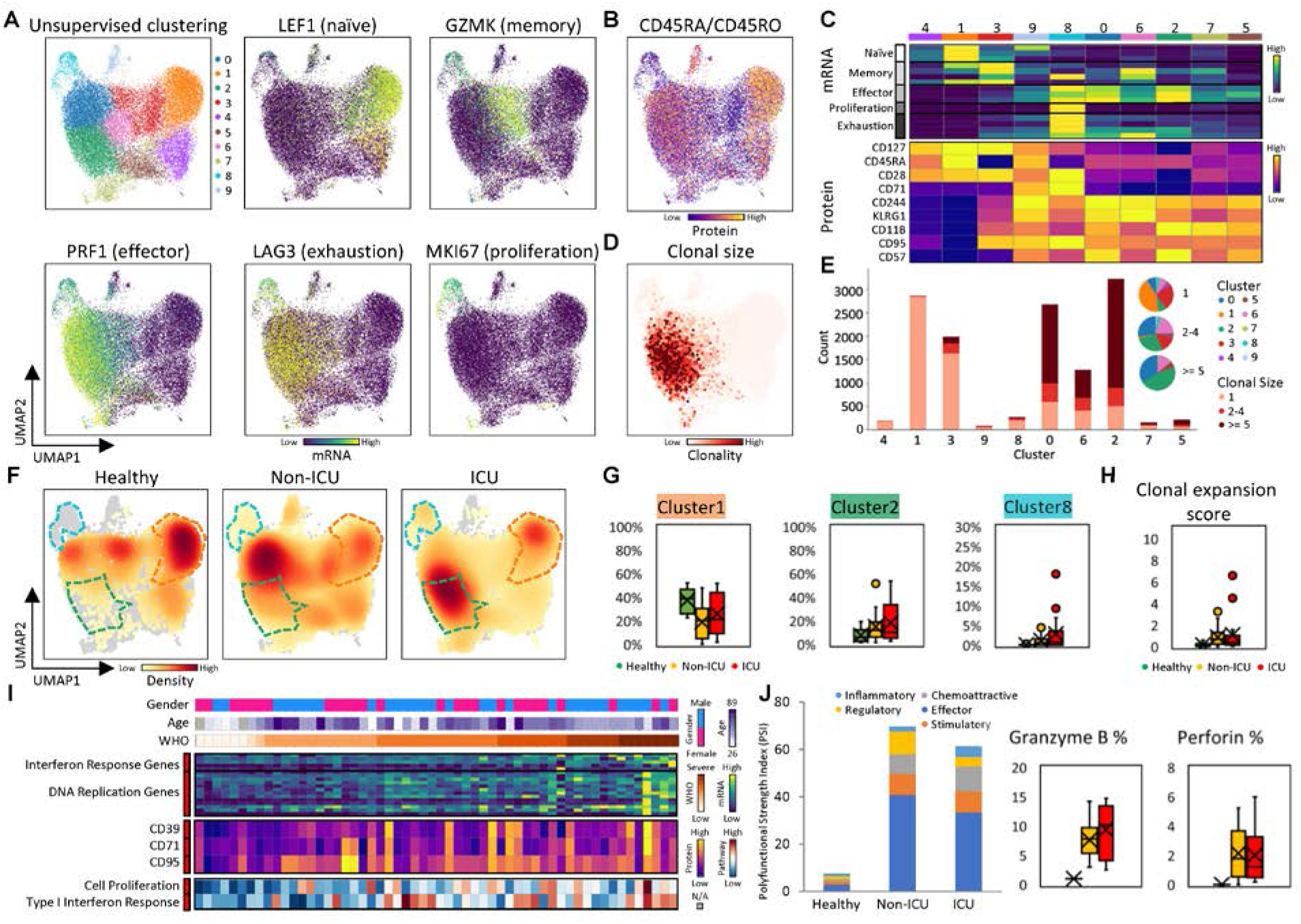
CD8+ T cell heterogeneity in COVID-19 patients and its association with disease severity. A. UMAP embedding of all CD8^+^ T cells colored by unsupervised clustering (top left panel) and other representative genes (other panels). B. UMAP embedding of CD8^+^ T cells colored by CD45RA/CD45RO surface protein ratio. C. Heatmaps showing the normalized expression of select genes (top panel) and proteins (bottom panel) in each cell cluster. D. UMAP embedding of CD8^+^ T cells shaded by T cell receptor clonal expansion level. E. Clonal expansion status, presented as bar plots, for each CD8^+^ T cell cluster, color-coded for clonal sizes of 1, 2-4, and >=5 clones. The pie charts shows the CD8^+^ T cell clonal composition percentage for each cluster. F. UMAP embedding of all CD8^+^ T cells for different blood draw samples (healthy, non-ICU, ICU). A few clusters that displayed significant density changes from group to group are encircled in dashed lines, colored according to the cluster identification from panel A. Orange, green and cyan dashed lines for clusters corresponds to naïve-like cell cluster 1, effector-like cell cluster 2, and proliferative CD8+ T cell cluster. G. Boxplots represent percentages of each highlighted CD8^+^ T cell cluster in F. over all CD8^+^ T cells for PBMCs from healthy donors (green), non-ICU blood draws (yellow), and ICU blood draws (red) of COVID-19 patients. H. Clonal expansion score, defined by ratio of cells with TCR clone size n>1 over cells with TCR clone size n=1, for CD8^+^ T cells in healthy donors (green), and non-ICU (yellow), and ICU (red) blood draws. I. Heatmap visualization of representative genes, surface proteins and pathways in CD8^+^ T cell that correlate strongly with disease severity, quantified by the WHO_ordinal_scale. Each column represents a sample and each row corresponds to levels of mRNA, surface protein, or GSEA pathway for the CD8+ T cells of that sample. Columns are ordered based on WHO ordinal scale in ascending order. Top three rows indicate the gender, age, and WHO ordinal scale. The heatmap keys are provided at right. Sidebar on the left of each row represent correlation of that value with WHO, with red (blue) indicating positive (negative) correlations. J. Functional characterization of CD8^+^ T cells using single-cell secretome analysis. Left panel: Single-cell polyfunctional strength index (PSI) of CD8^+^ T cells from healthy donors, and non-ICU, and ICU samples. The PSI (y-axis) is the numbers of polyfunctional cells in the sample, multiplied by the intensities of the secreted cytokines. Middle and right panel: Boxplots indicating the percentage of CD8^+^ T cells secreting GZMB and Perforin from samples from healthy donors (green), non-ICU blood draws (yellow), and ICU blood draws (red) of COVID-19 patients.

We next investigated the association of these subpopulations with disease severity. We projected the distribution density of cells from healthy, non-ICU and ICU COVID-19 samples onto the UMAP separately **(Figure 2F)**. As expected, the naïve-like cell cluster 1 **(orange dashed border in Figure 2F)** displayed the highest density in healthy patients. This subpopulation, along with naïve-T-cell-related genes and proteins, decreased in COVID-19 patients, **(Figures 2F, 2G and S2E)**. By contrast, effector-like cell cluster 2 **(green dashed border in Figure 2F)** and effector transcript for *GZMB* were enriched in COVID-19 patient samples relative to healthy donors **(Figures 2F, 2G and S2E)**. Consistent with this observation was that CD8^+^ T cell activation and differentiation markers (CD39, CD71, and CD95) exhibited positive correlations with disease severity, including the WHO ordinal scale **(Figure 2I)**. CD8^+^ T cell clonal expansion was also enhanced in COVID-19 patients **(Figure 2H)**. Furthermore, the proliferative CD8^+^ T cell cluster **(cluster 8, cyan dashed border in Figure 2F)** was enlarged in COVID-19 patients, especially in ICU samples **(Figures 2F and 2G)**. This observation was confirmed by the positive correlation of DNA replication transcripts and pathway enrichment score with disease severity **(Figure 2I)**. Thus, increased activation and clonal expansion of CD8^+^ T cells was present in the peripheral blood of COVID-19 patients, and the percentage of proliferative CD8^+^ T cells positively correlates with disease severity.

Notably, the most distinct and novel CD8^+^ T cell subpopulation is Cluster 8, which is nearly absent in healthy donors but elevated in ICU patients (**Fig 2G**). The dominant marker for cell proliferation (e.g. *MKI67)* is exclusively expressed in cluster 8, while, counterintuitively, this cluster also exhibits the highest levels of exhaustion markers, retains the 2nd highest cytotoxic signatures among all clusters, and has not fully lost the naïve signature (**Fig 2SF**). Such hybrid features of T cell exhaustion may be a unique signature of severe COVID19 infection.

Polyfunctional T cells produce multiple different cytokines and can release substantially higher amount of cytokines relative to other T cells, and so dominate the immune response to a pathogen relative to cells producing zero or one cytokine (Abel et al., 2010; Boyd et al., 2015; Lu et al., 2015; Ma et al., 2013; Parisi et al., 2020; Reyes-Sandoval et al., 2010; Zhou et al., 2017). To examine the polyfunctionality of T cells in SARS-CoV-2 infection, we applied single-cell secretome analysis (Isoplexis) to characterize the secretion of 32 cytokines by individual CD8^+^ T cells from COVID-19 patients characterized by different levels of disease severity. Polyfunctionality can be quantified by the Polyfunctional Strength Index (PSI), which is (the numbers of different proteins secreted) × (copy numbers secreted) (Lu et al., 2015; Ma et al., 2011). The PSI was up-regulated in all COVID-19 samples relative to healthy donors **(Figure 2J)**. In addition, the frequency of CD8^+^ T cells that secrete the cytotoxic granules granzyme B and perforin was also sharply increased **(Figure 2J)**. This is consistent with the relative increase of effector CD8^+^ T cells subpopulations in those patients **(Figure 2G)**. Thus, peripheral CD8^+^ T cells in hospitalized COVID-19 patients display increased polyfunctionality, along with an upregulated frequency of effector-cytokine-secreting T cells, which may be needed to control the virus infection in severe disease, yet may also contribute to cytokine release syndrome.

### CD4^+^ T cell proliferation, cytotoxic activity, and clonal expansion changes with severity of COVID-19

We next examined the association between disease severity and CD4^+^ T cell characteristics. Similar to the analysis of CD8^+^ T cells, we first assessed the subpopulation composition, and then interrogated the clonal expansion and polyfunctionality. The clustering displayed as color coded UMAP is shown in **Figure 3A** (top left panel). Naïve mRNA (*TCF7, CCR7*) and proteins (CD45RA and CD197) were uniquely upregulated in cluster 0 (blue) and 2 (green) **(Figures 3A, 3C, S3A and S3B)**. Th1 cytokine, *IFN-γ* and exhaustion markers (*LAG3*, CD279, and CD224) were all upregulated in cluster 3 (red) and cluster 8 (cyan) **(Figures 3A, 3C, S3A and S3B)**. Interestingly, CD244 (2B4) is also expressed in clusters 3 and 8, as well as cluster 7 (olive). This marker is more commonly seen in CD8^+^ T cells **(Figure 2C)** (Agresta et al., 2018)) and indicates the maintenance of an exhausted phenotype. Cluster 8 displayed uniquely upregulated proliferation genes *MKI67* and *TYMS* **(Figures 3A, 3C, and S3A)**. The Treg marker *FOXP3* was elevated mainly in cluster 1 **(Figures 3A and 3C)**. The CD45RA/CD45RO protein ratio was elevated in the naïve-like clusters 0 and 2 (**Figure 3B)**. Surprisingly, cluster 3 exhibited a strong elevation of the cytotoxic *PRF1* and *GNLY* cytokines **(Figures 3A, 3C and S3A)**. Cytotoxic CD4^+^ T cells can be induced by repeated antigen exposure (Juno et al., 2017; Takeuchi and Saito, 2017), suggesting that cluster 3 cells may have encountered viral antigens repeatedly. The high level of clonal expansion within cluster 3 **(Figures 3, 3E, S3C, and S3D)** further implies viral specificity of this cluster.

**Figure 3.**
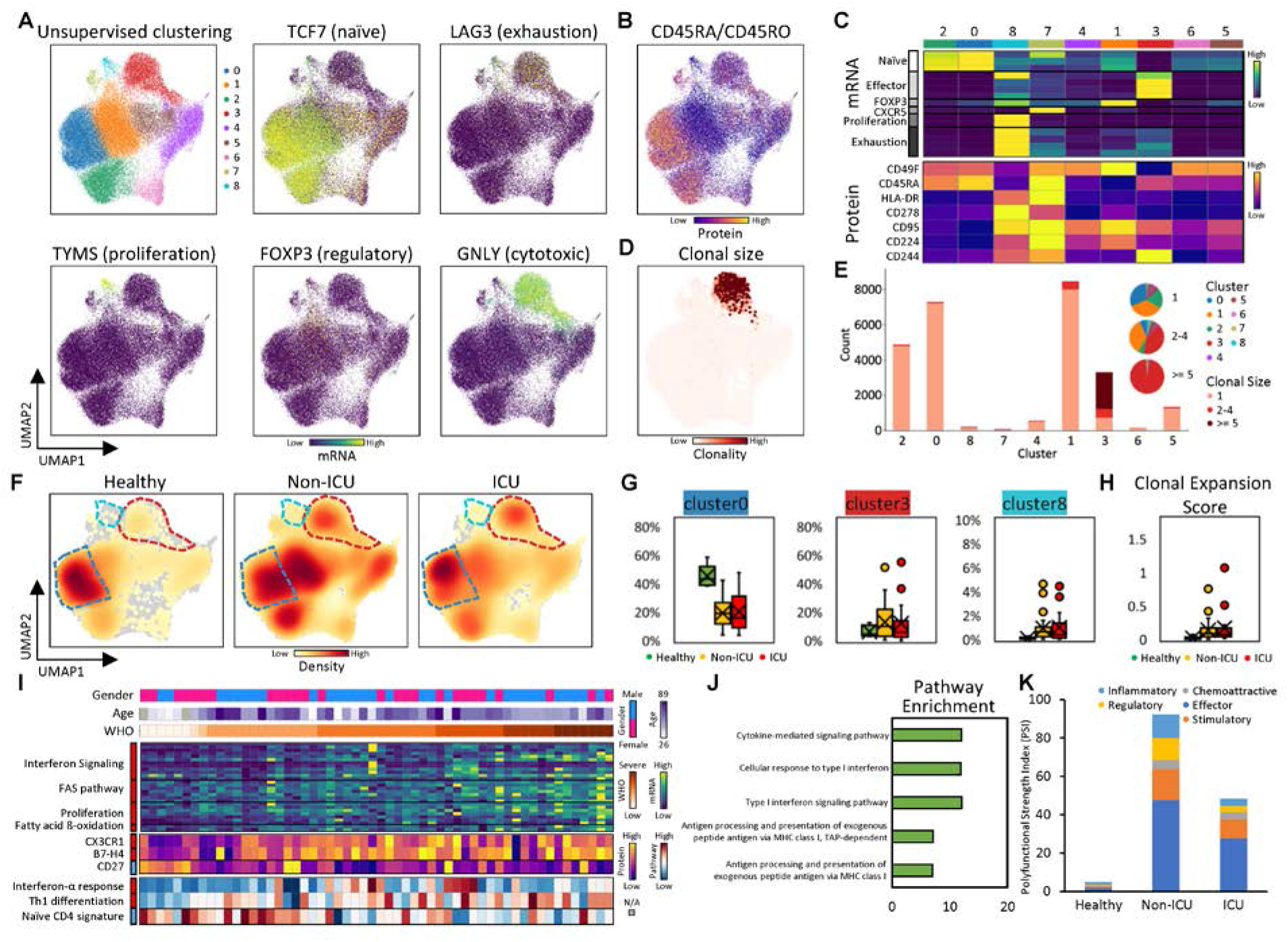
CD4^+^ T cell displayed heterogeneous proliferation, cytotoxic activity and clonal expansion levels and are associated with the severity of COVID-19. A. UMAP embedding of all CD4^+^ T cells colored by unsupervised clustering (top left panel) and other representative genes (other panels). B. UMAP embedding of CD4^+^ T cells colored by CD45RA/CD45RO surface protein ratio. C. Heatmap displaying normalized expression of select genes (top) and proteins (bottom) in each cell cluster identified in panel A. D. UMAP embedding of CD4^+^ T cells colored by clonal expansion level. E. Clonal expansion status, presented as bar plots, for each CD4^+^ T cell cluster, color-coded for clonal sizes of 1, 2-4, and >=5 clones. The pie charts shows the CD4^+^ T cell clonal composition percentages for each cluster. F. UMAP embedding of all CD4^+^ T cells for different blood draw samples (healthy, non-ICU, and ICU). A few clusters that displayed significant density changes from group to group are encircled in dashed lines, colored according to the cluster identification from panel A. Blue, red and cyan dashed lines corresponds to naïve like cluster (cluster 0), cytotoxic CD4^+^ T cell cluster (cluster 3), and proliferative cluster (cluster 8), respectively. G. Boxplots represent percentages of each highlighted CD4^+^ T cell cluster in F. over all CD4^+^ T cells healthy donors (green), non-ICU blood draws (yellow), and ICU blood draws (red) of COVID-19 patients. H. Clonal expansion score, defined by ratio of cells with TCR clone size n>1 over cells with TCR clone size n=1, for CD4^+^ T cells in healthy donors (green), non-ICU blood draws (yellow) and ICU blood draws (red) of COVID-19 patients. I. Heatmap visualization of representative genes, surface proteins and pathways in CD4+ T cell that correlate strongly with disease severity, quantified by the WHO_ordinal_scale. Each column represents a sample and each row corresponds to levels of mRNA, surface protein, or GSEA pathway for the CD4^+^ T cells of that sample Columns are ordered based on WHO ordinal scale in ascending order. Top three rows indicate the gender, age and WHO ordinal scale. The heatmap keys are provided at right. Sidebar on the left of each row represent correlation of that value with WHO, with red (blue) indicating positive (negative) correlations. J. Pathway enrichment analysis of genes that uniquely decrease in patients who improve (T2-T1, WHO decreased) in comparison with patients who did not. Each row represents a pathway. The x-axis represents the enrichment score defined by negative log10 P value. K. Functional characterization of CD4^+^ T cells using single-cell secretome analysis. The y-axis is the polyfunctional strength index (PSI) of CD4^+^ T cells from healthy donors, non-ICU blood draws, and ICU blood draws of COVID-19 patients.

CD4^+^ T cells from healthy, non-ICU and ICU COVID-19 samples are projected as density UMAP in **Figure 3F.** In comparison with healthy donors, the naïve like cluster (cluster 0, dark blue) was reduced in COVID-19 patients while the proliferative cluster (cluster 8, cyan) was increased **(Figures 3F and 3G)**. This result is consistent with the positive correlation between disease severity and proliferation related genes **(Figure 3I)**, and in line with a negative correlation between disease severity and the naïve cell pathway enrichment score **(Figure 3I)**. The novel cytotoxic cluster 3 cells are also upregulated in all COVID-19 patients **(Figures 3F and 3G)**, again possibly reflecting repeated SARS-CoV-2 viral antigen encounters for these T cell phenotypes. Such cytotoxic CD4^+^ T cells have, to our knowledge, not been reported before in COVID-19 patients. Finally, the proportion of clonally expanded CD4^+^ T cells was higher in COVID-19 patients **(Figure 3H)**, likely due to the expansion of virus-specific T cells to help control the infection.

We also quantified the genes, surface proteins, and pathway enrichment scores in CD4^+^ T cells and their correlation with disease severity. Consistent with recent literature (Chen et al., 2020; Mathew et al., 2020), an increase of FAS pathway genes, proliferation genes, activation/effector surface markers CX3CR1, B7-H4, the Th1 differentiation pathway score, and the Type I interferon response gene signature were positively correlated with disease severity **(Figure 3I)**. By contrast, naïve T cell signatures and the early-activation marker CD27 were negatively correlated **(Figure 3I)**.

As blood draws from two different time points were taken for the same patient, we also analyzed individual patient data for time-progression trajectories to discern gene expression dynamics and their associations with alleviated or aggravated disease symptoms. Type I interferon response genes, cytokine signaling activities, and antigen presentation gene signature all decrease in patients who improve from T1 to T2 (WHO score decreases) **(Figure 3J, Table S7)**. Thus, the Type I interferon response, T cell activation and differentiation processes strongly associate with disease severity and its progression over time.

We analyzed for the polyfunctionality of single, viable CD4^+^ T cells using the PSI (Lu et al., 2015; Ma et al., 2011). We found that CD4^+^ T cells from COVID-19 patients demonstrated a greatly elevated polyfunctionality relative to those from healthy donors **(Figure 3K)**. Within COVID-19 patients, CD4^+^ T cells from ICU samples exhibited a decrease in polyfunctionality **(Figure 3K)**. Specifically, the contributions from effector cytokines, stimulatory cytokines, and inflammatory cytokines (as defined in the Methods) were all reduced in ICU samples **(Figures 3F and S3F)**. The decrease of polyfunctionality in ICU samples may be a signature of the dysregulated immune function in the most severely infected patients. Notably, a similar decrease in the PSI of CD8^+^ T cells was also observed for ICU samples **(Figure 2J)**. This may be a general trend reflecting T cell exhaustion in the most critically ill patients.

### B cell differentiation status into plasmablasts increases with COVID-19 disease severity

B cells are the other major player of the adaptive immune system, and stimulating them to produce virus neutralizing antibodies is a major objective of current vaccine efforts. Utilizing unsupervised clustering, B cells were partitioned into three major phenotypes **(Figures 4A, 4B, S4A, and S4B)**: naïve B cells (IgD^+^IgM^+^CD27^-^), memory B cells (CD27^+^CD38^low^CD24^high^) and plasmablasts/plasma cells (CD20^-^ CD38^high^CD27^+^). The proportions of all three phenotypes displayed an obvious departure progressively from healthy to COVID-19 non-ICU to ICU samples. Naïve cell (cluster 0 (blue)) percentages and gene signatures were decreased in patients with higher disease severity, whereas plasmablast/plasma cell (cluster 4 (purple)) frequencies and gene signatures were elevated with more severe infection **(Figures 4A,4E–4H, and S4C–4E)**. Memory B cell (cluster 2 (green)) percentages increased in COVID-19 patients compared to healthy donors. Class-switched memory B cells that expressed IgA and IgG genes *(IGHG1* and *IGHA1)* represented the major cell population of memory B cells identified **(Figures 4A and S4A)**. Meanwhile, the levels of naïve B cell surface proteins, IgD and IgM, were negatively correlated with disease severity **(Figures 4H and S4D)**. B cell activation signatures including downregulation of *FCER2*, and upregulation of *SLAMF7* and CD11C, were more prominent in COVID-19 patients compared to healthy controls **(Figures 4G, S4C and S4D)**. Transcripts of antibody-secreting cell (ASC)-related genes and surface markers, including CD138, *XBP1, SPCS3*, and *IGHG4*, were over-represented in COVID-19 patient samples **(Figures 4H and S4C)**. Indeed, marked plasmablast expansion has been associated with disease severity in infectious diseases such as acute dengue virus infection (St. John and Rathore, 2019). We also found pronounced IgA *(IGHA1)* gene expression in both COVID-19 non-ICU and ICU samples compared to healthy samples, with the highest expression levels found in ICU samples **(Figure S4C)**, suggesting a role for the IgA response in mucosal immune responses targeting viral control. Additionally, as seen in monocytes, MHC-class II genes in B cells were also downregulated in COVID-19 patients in comparison with healthy donors, and such downregulation was more pronounced in ICU samples compared to non-ICU samples **(Figures 4G and S4C)**, as previously reported (Wilk et al., 2020). Such altered MHC class II expression could potentially permit evasion of immune recognition of viral epitopes by CD4^+^ T cells, thus impairing the adaptive immune response in COVID-19 patients. Overall, our findings suggest that higher levels of B cell activation and differentiation into ASCs is associated with more severe disease.

**Figure 4.**
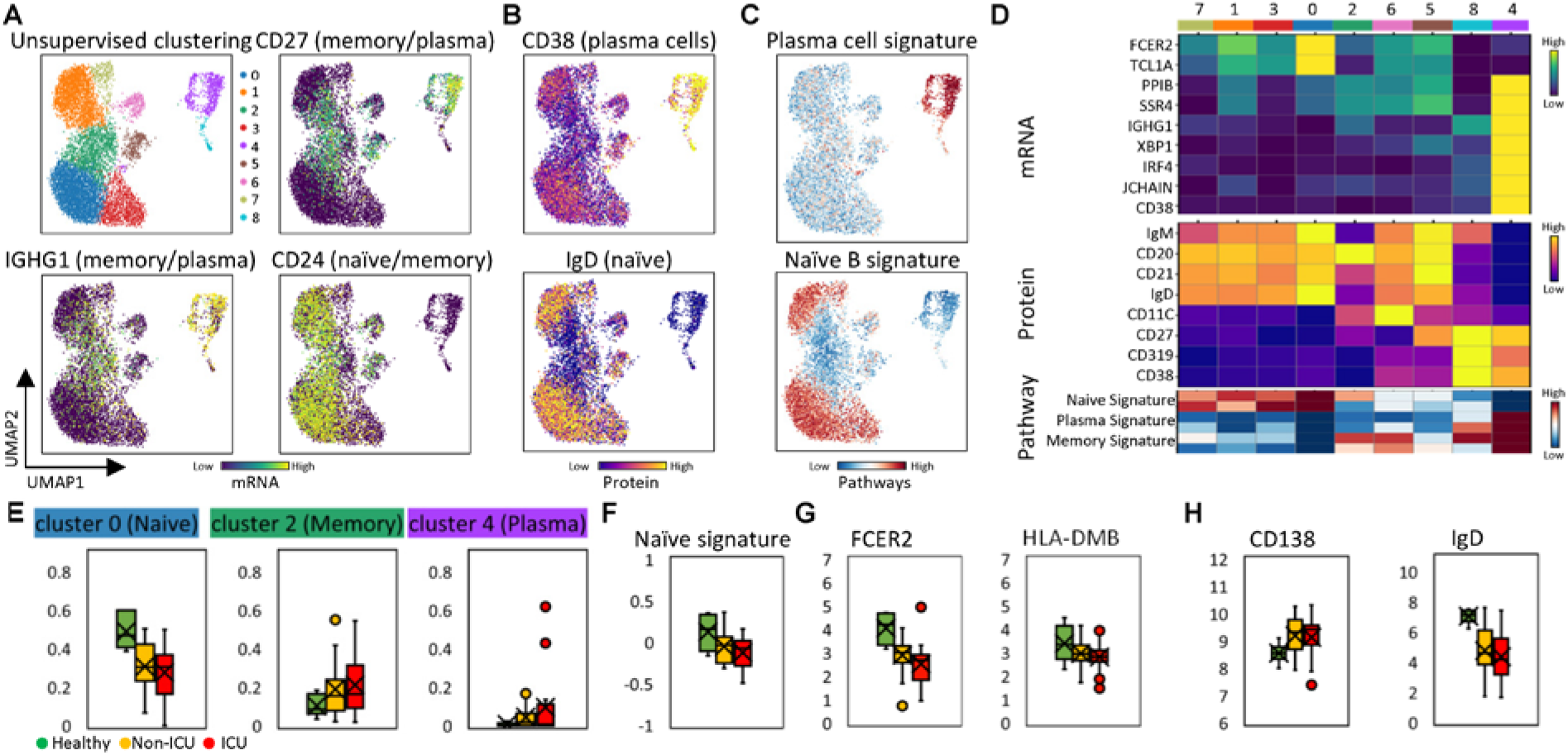
Deep profiling of COVID-19 patient B cell populations reveals heterogeneous differentiation status that correlates with disease severity. A. UMAP embedding of all B cells colored by unsupervised clustering (top left panel) and other representative genes (other panels). B. UMAP embedding of all B cells colored by levels of two representative surface proteins. C. UMAP embedding of all B cells colored by representative pathway enrichment scores. D. Heatmap displaying normalized expression of select genes (top) and proteins (middle) and pathway signatures (bottom) in each B cell cluster. E. Boxplots indicating percentage of B cells in select clusters, from samples collected from healthy donors (green), non-ICU blood draws (yellow), and ICU blood draws (red) of COVID-19 patients. F-H. Boxplots showing the mRNA expression levels (panel F), surface protein levels (panel G), and the levels of pathway enrichment scores (panel H) in healthy donors (green), non-ICU blood draws (yellow), and ICU blood draws (red) of COVID-19 patients.

### Decreased antigen presentation and increased extravasation and polyfunctional potential in monocytes are associated with COVID-19 disease severity

The relative frequency of monocytes appeared to markedly increase in COVID-19 patients compared to healthy donors **(Figure 1F)**. UMAP visualization of all monocytes showed a clear separation of non-classical monocytes (CD14^low^ CD16^+^ cluster 3, red), characterized by the upregulated genes *FCGR3A, CX3CR1* and surface proteins CD16 **(Figures 5A, 5B, 5D, and S5A)**. Among the classical monocyte clusters (CD14^+^ CD16^-^), cluster 1 (orange) was distinguished by increased expression of the extravasation marker *CCR2*, type I interferon (IFN) response genes, and inflammation related genes such as *S100A9, S100A12*, and a decreased antigen stimulus response gene signature **(Figures 5A, 5C, 5D, S5A, and S5B)**. We examined the association of these subpopulations with disease severity. Proinflammatory cluster 1 classical monocytes were more prevalent in ICU samples **(Figure 5E)**. Accordingly, the cluster1-unique markers, proinflammatory gene *S100A8*, and extravasation gene *CCR2*, were upregulated in ICU samples **(Figure 5F)**. This is consistent with the observation that extravasation genes, monocyte activation, and activationassociate metabolic pathways were positively correlated with disease severity **(Figure 5H)**. Accordingly, upon stimulation with LPS, the frequency of monocytes that secrete proinflammatory cytokines IL-6, IL-8, etc., also increased in COVID-19 patients, especially in ICU samples **(Figure S5C)**. Compared to cluster 1, the proportion of non-classical monocytes (cluster 3), which are generally better at antigen presentation and restoring homeostasis (Schmidl et al., 2014), displayed a sharp decrease in ICU samples **(Figure 5E)**. HLA class II genes, proteins, and antigen presentation pathway enrichment scores were also down-regulated, especially in ICU samples **(Figures 5F and 5G)**. In fact, HLA-class II gene expression also showed a strong negative correlation with disease severity **(Figure 5H)**. This is reminiscent of monocyte “immunoparalysis” reported in sepsis where, under conditions of high inflammation, the monocytes become dysfunctional (Giamarellos-Bourboulis et al., 2020). This is further supported by the decrease in *TNF-α* transcripts **(Figure 5F)**. In fact, the expression of MHC class II genes across COVID-19 samples is negatively correlated with IL-6 plasma level **(Figures 5I and S5D)**, indicating that monocyte dysfunction is likely related to and possibly induced by the hyper-inflammatory plasma environment.

**Figure 5.**
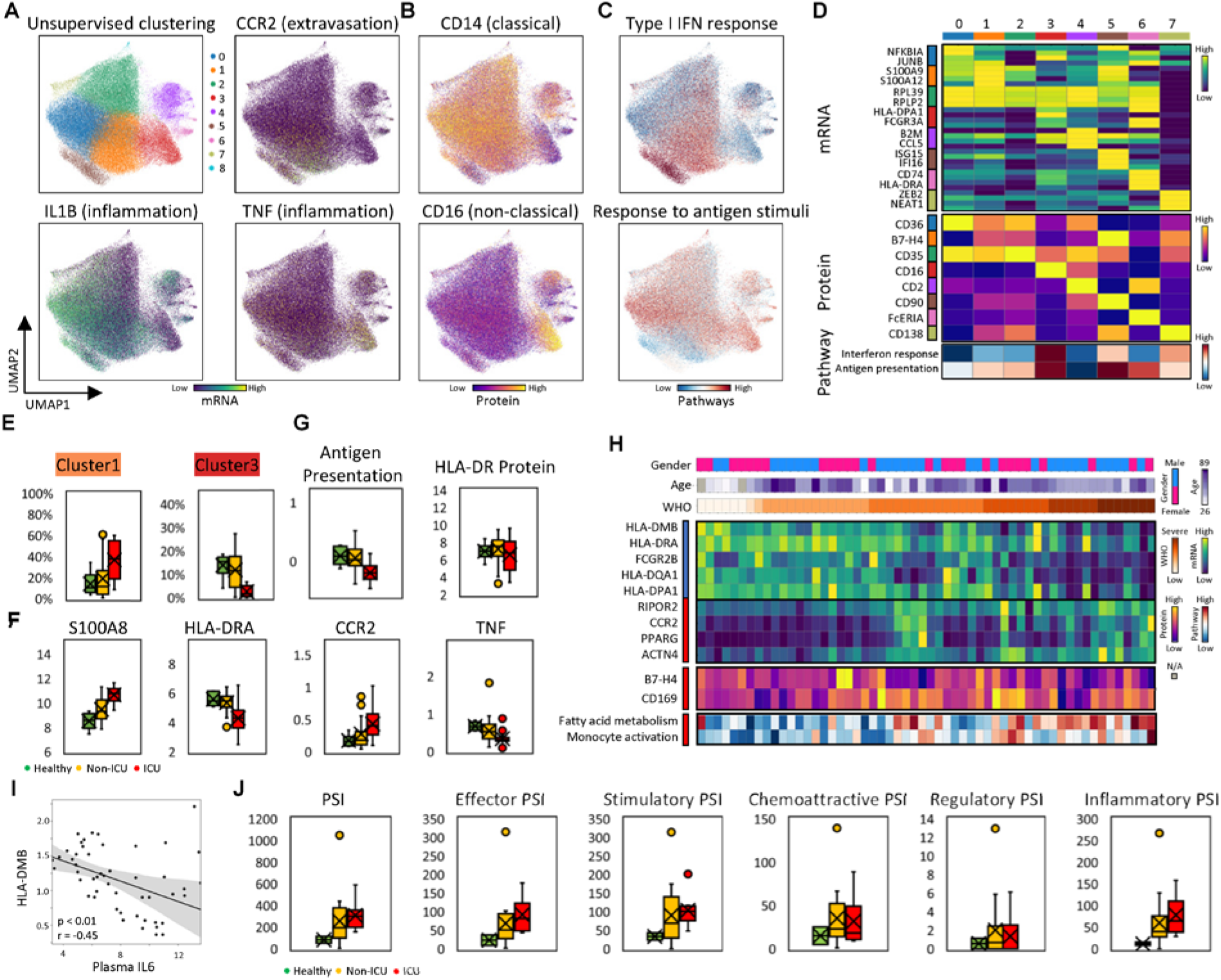
Alterations of antigen presentation and inflammation in monocytes are associated with COVID-19 disease severity. A. UMAP embedding of all monocytes colored by unsupervised clustering (top left panel) and other representative genes (other panels). B. UMAP embedding colored by the levels of protein markers for classical (CD14) and non-classical (CD16) monocytes. C. UMAP embedding of monocytes colored by representative pathway enrichment scores. D. Heatmap displaying normalized expression of select genes (top) and proteins (middle) and pathway signatures (bottom) in each monocyte cell cluster. E. Boxplots indicating cell percentages in select monocyte clusters from samples in healthy donors (green), non-ICU blood draws (yellow), and ICU blood draws (red) of COVID-19 patients. F. Boxplots showing the levels of four representative mRNA from monocytes in healthy donors (green), non-ICU blood draws (yellow), and ICU blood draws (red) of COVID-19 patients. G. Boxplots showing the pathway enrichment score (left panel) and surface protein levels (right panel) from monocytes in healthy donors (green), non-ICU blood draws (yellow), and ICU blood draws (red) of COVID-19 patients. H. Heatmap visualization of representative genes, surface proteins and pathways in monocytes that correlate strongly with disease severity quantified by the WHO ordinal scale. Each column represents a sample and each row corresponds to levels of mRNA, surface protein, or GSEA pathway for the monocytes of that sample. Columns are ordered based on WHO ordinal scale in ascending order. The top three rows indicate the gender, age, and WHO ordinal scale. The heatmap keys are provided at right. Sidebar on the left of each row represent correlation of that value with WHO, with red (blue) indicating positive (negative) correlations. I. Spearman correlation of HLA-DMB gene expression within monocytes with plasma IL6 level. The regression line is indicated in black, with a 95% confidence interval shown in shaded gray. Spearman Correlation coefficient and associated p-value shown. J. Functional characterization of monocytes from PBMC samples using single-cell secretome analysis. Single-cell polyfunctional strength index (PSI) of monocytes from healthy donors, non-ICU blood draws, and ICU blood draws of COVID-19 patients. Other plots are sub-category specific PSI, which is calculated by only considering those cytokines representing each sub-category, as indicated in the box plot titles.

We also investigated polyfunctionality by assaying for the PSI of monocytes collected from healthy, non-ICU and ICU samples after LPS stimulation *ex vivo*. In comparison with monocytes from healthy donors, elevated functionality was observed in stimulated monocytes from COVID-19 donors, especially in those ICU samples **(Figure 5J)**. The polyfunctionality contributions from effector, stimulatory and inflammatory cytokines all increased with disease severity **(Figure 5J)**.

In summary, reduced antigen presentation and enhanced extravasation and polyfunctional potential in monocytes are associated with disease severity in COVID-19 patients.

### Proliferation and maturation in NK cells vary with COVID-19 disease severity

As another important player in controlling virus infection, NK cells were also analyzed at single cell resolution. UMAP visualization of clustering showed a separation of two major subsets, which corresponded to the two well-known NK cell subsets: CD56^bright^ (cluster 4 (purple)) and CD56^dim^ CD16^bright^ (cluster 0-3 (blue, orange, green, and red) and 5-6 (brown and pink)) NK cell subpopulations **(Figures 6A, 6B, and 6D)**. As expected, the CD16^bright^ subpopulation displayed high levels of cytotoxic transcripts **(Figures 6A, 6B, and 6C)**. Interestingly, among CD56^dim^ CD16^bright^ NK cells, one subset (cluster 5) exhibited a pronounced type I IFN response gene signature **(Figures 6A, 6C and 6D)**, and the other subset (cluster 6) showed the highest level of the proliferation marker *MKI67* among all the NK cell subsets **(Figures 6A, and 6D)**. The percentage of proliferative cluster 6 NK cells showed a pronounced increase in ICU samples compared to non-ICU samples and healthy controls **(Figure 6E)**. Consistently, DNA replication and cell proliferation associated genes and pathways were also positively correlated with the WHO disease severity score **(Figures 6F and 6G)**. Different from cluster 6, the percentages of the CD56^bright^ NK cell subpopulation (cluster 4) and its representative marker *SELL* decreased in COVID-19 patients, especially in non-ICU samples, compared to healthy donors **(Figures 6E, and 6F)**, which may reflect the skewing of NK cells from cluster 4 to more cytotoxic phenotypes.

**Figure 6.**
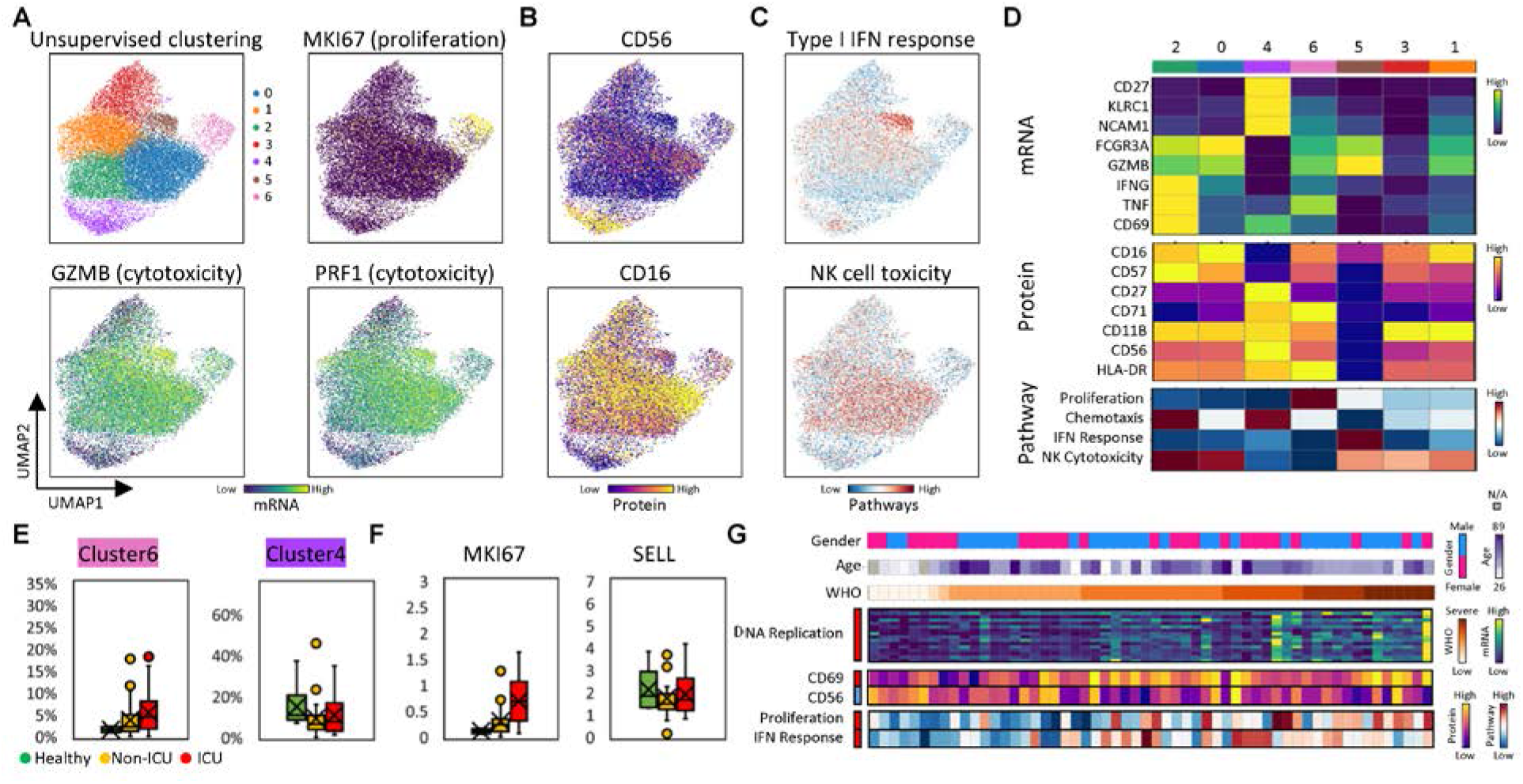
Deep profiling of COVID-19 patient NK cell populations reveals heterogeneous proliferation that correlates with disease severity. A. UMAP embedding of all NK cells colored by unsupervised clustering (top left panel) and other representative genes (other panels). B. UMAP embedding of all NK cells colored by the levels of two representative surface proteins. C. UMAP embedding of all NK cells colored by representative pathway enrichment scores. D. Heatmap displaying normalized expression of select genes (top) and proteins (middle) and pathway signatures (bottom) in each NK cell cluster. E. Boxplots indicating percentage of select NK cell clusters from samples in healthy donors (green), non-ICU blood draws (yellow), and ICU blood draws (red) of COVID-19 patients. F. Boxplots showing the levels of two representative transcripts from NK cells in healthy donors (green), non-ICU blood draws (yellow), and ICU blood draws (red) of COVID-19 patients. G. Heatmap visualization of representative genes, surface proteins, and pathways in NK cells that correlate strongly with disease severity, as quantified by the WHO ordinal scale. Each column represents a sample and each row corresponds to levels of mRNA, surface protein, or GSEA pathway for the NK cells of that sample. Columns are ordered based on WHO ordinal scale in ascending order. The top three rows indicate the gender, age, and WHO ordinal scale. The heatmap keys are provided at right. Sidebar on the left of each row represent correlation of that value with WHO, with red (blue) indicating positive (negative) correlations.

We also observed correlations between CD69, a type I IFN inducible marker, and genes associated with IFN responses with disease severity **(Figure 6G)**. This may be associated with the elevated inflammatory environment in the blood or in tissues **(Figure 1C)**. The pronounced proliferation and type I IFN responses of NK cells indicated their vigorous activation upon SARS-CoV-2 infection. We also viewed each patient as a time-progressing trajectory and examined the changes of gene expression and their association with alleviated or aggravated disease symptoms. We found that expression of cytotoxic NK cell marker CD16 and terminal differentiation marker CD57 are both decreased in patients who improved (decrease of WHO from T1 to T2) versus those that did not **(Figure S6D, Table S7)**. This indicates that reduced levels of cytotoxic, terminally differentiated NK cells are associated with COVID-19 recovery. Thus, NK cell proliferation and maturation status also reflect the severity of SARS-CoV-2 infection.

### Integrating multi-omic profiles from different cell types resolves an orchestrated response gene module of innate and adaptive immune cells that correlates with disease severity

As successful control of virus infection requires precise cooperation between both the innate and the adaptive immune cells, we hypothesized that the coordinated changes of multi-omic features across different cell types could provide a more unified view of the immune response in COVID-19 patients. A major challenge towards capturing such coordination is the huge number of genes expressed in different cell types coupled with inter- and intra-patient heterogeneity and associated variations in COVID-19 infection severity. We hypothesized that many genes co-vary across samples and thus constitute functional gene modules. In other words, genes from different cell types may co-vary and therefore reside in the same gene module, thus providing a simplifying insight into how different cell types coordinate their behaviors. To capture such coordinated changes, we utilized Surprisal Analysis (Remacle et al., 2010; Zadran et al., 2014), which has been applied to consolidate multi-omics bulk and single-cell data (Kravchenko-Balasha et al., 2014, 2016; Remacle et al., 2010; Su et al., 2017, 2019, 2020). The goal is to condense the changes of millions of correlated genes from different cell types into changes in just one major gene module or axis of expression, which contains the co-varying genes from all major immune cell types. As a result, each sample is reduced into a single dot along that gene module axis **(Figure 7A)**. This data integration could permit the delineation of correlated changes and disease stages.

**Figure 7.**
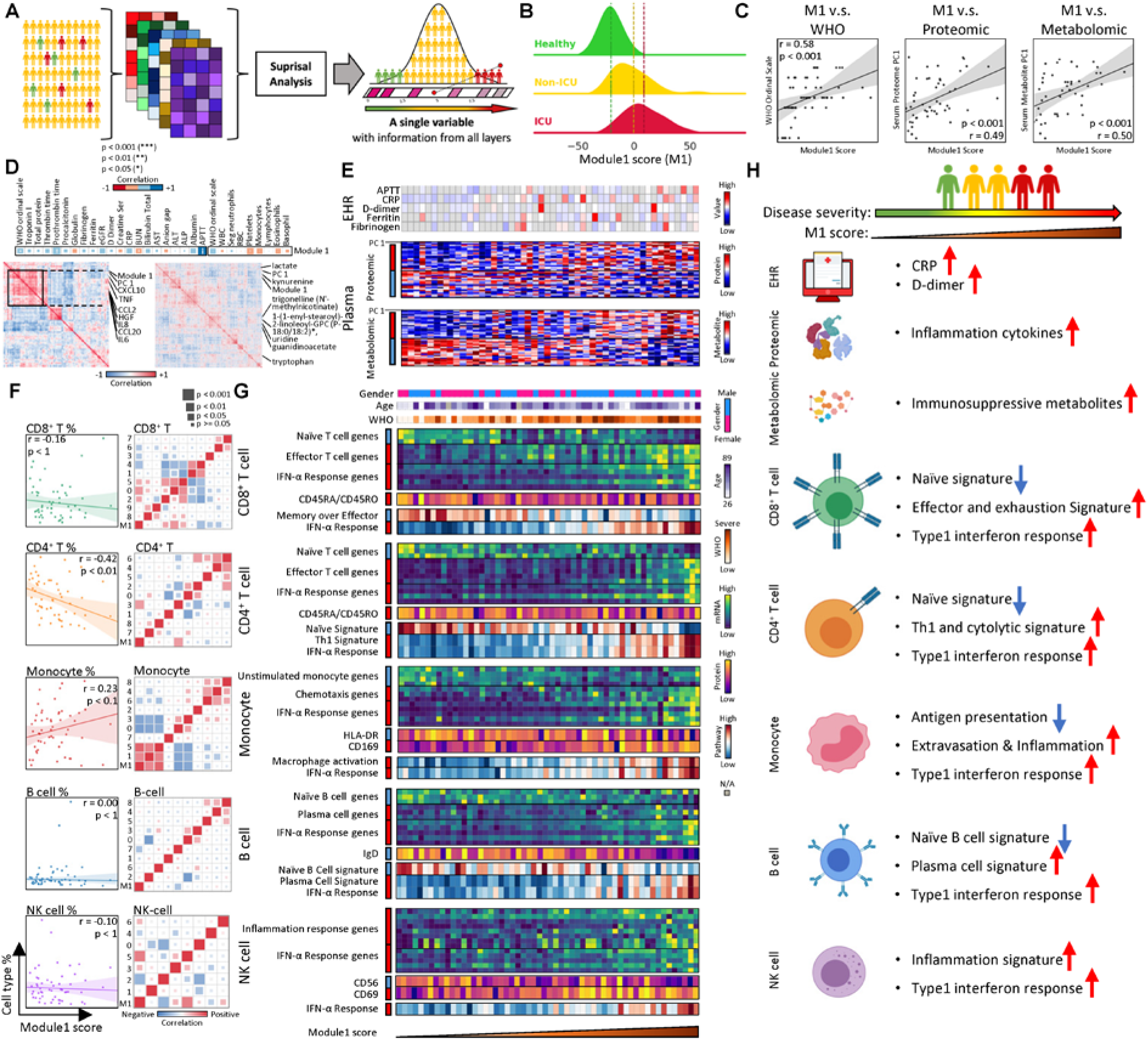
Integrating multi-omic profiles across immune cell types resolves an orchestrated gene module that correlates with disease severity. A. Illustration of integrating data from different immune cell types from all samples, followed by reduction in single a dimensional representation (gene module 1 (M1)) using surprisal analysis. B. Distribution of individual PBMC data sets along M1 for healthy donors (green), non-ICU (yellow), and ICU (red) patients. C. Spearman correlations of M1 with disease severity, as measured by the WHO ordinal scale), and with principal component (PC)1 of the plasma proteomic data and PC1 of the plasma metabolomics data (in Figure 1C and 1D). Regression lines are indicated in black, with 95% confidence area in shaded gray. Spearman Correlation coefficient and associated p-value shown. D. Correlations between M1 with clinical data (top panel: 1-dimensional map) and plasma proteomic and metabolomics data (bottom left and right panels, respectively). For the clinical data, the square size corresponds to absolute value of the correlation coefficient. Blue indicates positive correlation and red indicate negative correlation (*p < 0.05, **p < 0.01, ***p < 0.001). E. Heatmap visualization of EHR clinical data, plasma proteomic and metabolomic data that correlates with gene module1. Each row corresponding levels of clinical data, blood protein, or metabolite of that sample and each column represent a blood draw sample. Columns are ordered based on M1 score in ascending order. Sidebar on the left of each row represent correlation of that value with M1 score, with red (blue) indicating positive (negative) correlations. F. Correlation of immune cell type percentage and subtype percentages with M1 score. Left panel: Spearman correlations of M1 with percentage of five immune cell types. Right panel: Correlation matrix for percentage of different subtypes (based on previous clustering) within the five immune cell types and M1. G. Heatmap visualization of representative genes, surface proteins and pathways for each cell type that correlate positively or negatively with M1. Each column represent a PBMC sample and each row represents measured levels of transcripts, surface proteins, or pathways enrichment score within each cell type (indicated on the left) for that PBMC sample. Columns are ordered based on M1 score in ascending order. Top three rows indicating the gender, age, and WHO ordinal scale for each sample. Keys are presented in the sidebars at right. Sidebar on the left of each row represents the marker’s correlation with the M1 score, with red (blue) indicating positive (negative) correlations. H. Summary for how the clinical data, the plasma proteomic and metabolomics profiles, and how the major immune cell subtypes and their associated feature change along M1 axis. Red up (blue down) arrows represent positive (negative) correlations with M1 score.

The dominant gene module, module1 (M1), which was computed only from immune cell behavior without considering any clinical information, readily separates samples from healthy, non-ICU and ICU samples **(Figure 7B)**. The M1 score of each sample also strongly correlated with disease severity **(Figure 7C, Spearman Correlation coefficient = 0.58, p < 0.001)**. Furthermore, the M1 score, which was calculated without considering the multi-omic plasma data **(Figure 1C)**, also independently showed a strong correlation with the plasma proteomic principal component (PC)1 and metabolomics PC1 **(Figure 7C)**, both of which are aligned with disease severity **(Figure 1C)**. These results indicate that coordinated immune response across cell types can be captured by a single gene module and is associated with the severity of the disease.

We queried for clinical features that correlate with M1. We found that CRP, D-dimer and ferritin, which were upregulated in ICU samples, positively correlate with M1 **(Figures 7E and S7C)**, indicating the patients with a higher M1 score likely exhibit elevated levels of these clinical features. Among plasma proteins and metabolites analyzed, many proinflammatory cytokines, including IL6, TNF, CXCL10 and CCL20 etc., were all positively correlated **(Figures 7D, 7E and S7A)**. Tryptophan was negatively correlated with M1, and its downstream immunosuppressive metabolite kynurenine exhibited a positive correlation **(Figures 7D, 7E, and S7B)**. These results indicate that immune cell behaviors, reflected in M1, can be associated with the proinflammatory proteomic and immunosuppressive metabolic environment in the plasma.

We further examined the detailed features of adaptive immune systems and their association with module1 increase. As we demonstrated independently, the percentages of CD4^+^ and CD8^+^ T cell were both negatively correlated with M1, while the monocyte fraction showed a positive correlation **(Figure 7F)**. Within each major immune cell type, different subpopulations exhibited differential correlations with M1. In CD8^+^ and CD4^+^ T cells, effector-like cytotoxic cell clusters (cluster 0 and 2 for CD8^+^ T cell, cluster 3 for CD4^+^ T cell) positively correlated with M1 **(Figure 7F)**. Consistently, cytotoxic genes, effector proteins and pathway signatures also positively correlated with the M1 score. In contrast, naïve and memory like features were negatively correlated **(Figures 7G and S7D)**. In B cells, naïve cell cluster 7 showed a negative correlation **(Figure 7F)**, which is consistent with the switching from a naïve to a plasma cell signature with increasing M1 **(Figure 7G)**.

Innate immune cells also showed transcriptional alterations along the M1 axis. In monocytes, the proinflammatory cluster 1 showed a positive correlation and the non-classical monocyte cluster 3 showed a negative correlation. Accordingly, increased pro-inflammatory gene signatures and decreased MHC class II genes in monocytes associates with an increase in the M1 score **(Figures 7G and S7D)**. The NK cell inflammation response genes also positively correlate with M1. Meanwhile, as M1 increases, cytotoxic and proliferative NK cells also increase **(Figure 7G)**, with increases in inflammatory response genes. Importantly, we show that type I IFN-response genes and related pathways exhibit a positive correlation with M1 across all immune cell types **(Figure 7G and S7D)**, suggesting that the immune response may be dominated by the type I IFN responses. This has been reported to be upregulated in virus infection (Murira and Lamarre, 2016; PERRY et al., 2005; Stetson and Medzhitov, 2006), further confirming the essential role of type I IFN response in COVID-19 immune response and providing confidence to this analysis.

In summary, by integrating deep profiling of all major immune cell types across samples, we integrated the immune response from all cell types (with more than a million variables) and condensed all of them into a single gene module **(Figure 7A)**. This gene module is associated with activated IFN responses across all major immune cell types and reflects cell-type specific differentiation and activation processes. This immune response gene module also independently correlates with the plasma multi-omic analytics, specific clinical observations, and measured disease severity. This result suggests that coordinated immune responses across innate and adaptive immune cell types are strongly linked to disease severity and clinical outcome. It also suggests that certain clinical observations may be interpreted as surrogates for assessing these immune responses, but such validation will require the analysis of larger patient cohorts.

## DISCUSSION

A deep understanding of patient immune responses to COVID-19 is fundamental to defining the effectiveness of patient treatments, predicting disease prognosis, and for understanding the reported heterogeneity of infection severities. We have provided a comprehensive view of immune status sampled through serial blood draws over an approximately 1-week period following diagnosis and hospitalization for 26 COVID-19 patients. Time-dependent clinical trajectories were integrated with comprehensive probes of plasma proteomics and metabolomics, and multi-omic single cell analytics of five major immune cell classes.

There are a number of novel observations for which there is not large literature precedence and may provide important clinical insights. For example, T cell exhaustion has been frequently reported in severe SARS-CoV2 infections (Diao et al., 2020). Indeed, transcriptomic analysis of our CD8^+^ T cells reveals elevated exhaustion markers for multiple clusters in the UMAP of **(Figure 2A).** However, cluster 8, which exhibits the highest exhaustion marker levels simultaneously exhibits the highest proliferation signature **(Figure S2F)**. This finding is counterintuitive as exhausted T cells are reported with a marked absence of proliferation markers (Saeidi et al., 2018). In fact, the co-occurrence of these two features is also observed in CD4^+^ T cells **(cluster 8 in Figure S3A and S3B).** Interestingly, the most clonally expanded CD4^+^ T cell populations are not in this exhausted cluster but in the nearby cluster 3 comprised of equally novel, cytotoxic CD4^+^ T cells **(cluster 3 in Figure S3A)**. Thus, the cluster 8 exhausted and proliferative CD4^+^ T cells may not be virus specific. The presence of these novel CD8^+^ and CD4^+^ subpopulations should be considered regarding the use of T cell activating immune-checkpoint inhibitors in COVID-19 treatments.

Immunosuppressive agents, such as tocilizumab and steroids, are also being used to treat COVID-19 patients to mitigate the hyperinflammatory state in COVID-19 although such immunopathology is still debated (Vardhana and Wolchok, 2020). Monocyte immunoparalysis (Lukaszewicz et al., 2009) has also been considered as a protective immune-response towards hyperinflammation. It is characterized by a robust downregulation of MHC-class II and TNF transcripts, and has provided a robust prognosis marker for fatal outcomes in sepsis (van Ton et al., 2018). Thus, its occurrence in COVID-19 patients, especially for those in the ICU, implies similarities between these two conditions. Our single-cell monocyte transcriptomic and plasma proteomic data also reveals a strong anticorrelation between plasma IL6 and HLA class II transcripts in monocytes, suggesting a causal inhibitory relationship between the two, which has been validated in other biological contexts (Ohno et al., 2016). This also implies that the resultant decrease of MHC-class II transcripts in monocytes may impair adaptive T cell responses.

In addition to the aforementioned novel observations from investigating each cell type in isolation, the human immune response to viruses should also be viewed as a finely tuned biological orchestra, in which different cell types need to coordinate with one another in harmony to ensure proper viral clearance. In fact, a major insight from our approach is the reduction of over 1 million multi-omic immune-features into a single axis gene module M1, which provides an orchestrated view of the immune response to SARS-CoV-2 that can be directly correlated with clinical features. For example, our analysis showed strong correlations between M1, the WHO severity score, and many clinical features including C-reactive protein (poor prognosis marker for COVID19 (Yan et al., 2020)), and the blood coagulation biomarkers prothrombin time (PT) and activated partial thromboplastin time (APTT), providing immune correlates of the well-reported blood clotting in COVID-19 patients. This simplified gene module can also significantly simplify future evaluations of COVID-19 patients as their immune response could be predicted by a few representative markers associated with M1. For example, Type I IFN response genes, whose transcript levels universally correlate with M1 across all five major immune cell types, may serve as valuable biomarkers for predicting patients’ disease severity and various clinical features (e.g. cytokine release syndrome etc.).

Further observations point to the value of the integrated approach described here. For example the association of COVID-19 infection severity with the elevation of kynurenine and the drop in its molecular precursor, tryptophan (**Figure 1D**), suggests activation of indoleamine 2,3-dioxygenase (IDO) (Wang et al., 2010). IDO typically is expressed at very low levels, but, in vascular endothelial cells (Hansen et al., 2000), it can be induced by IFN-γ, which we also find elevated with increasing disease severity (**Figure 1C**). Kynurenine is a vascular relaxing factor that, when elevated, would be expected to manifest as hypotension (Wang et al., 2010). In fact, hypotension has indeed been reported as a dominant characteristic of critically ill COVID-19 patients (Bhatraju et al., 2020). A primary source of IFN-γ is often Th1 CD4^+^ lymphocytes. Those cells, which are located in cluster 3 of **Figure 3A** (and **Figure S3A**), are the ones that exhibit the novel cytotoxic function and are uniquely populated by those CD4^+^ T cells that have clonally expanded (**Figures 3D, 3E and S3C**). Th1 differentiation of CD4^+^ T cells is also found to increase with infection severity (**Figures 3I and 7G**), while naïve CD4^+^ T cell genes are reduced. Notably, this Th1 gene signature in CD4^+^ T cells closely tracks the Type I IFN response that is seen across all cell types and significantly correlates with M1 (**Figure 7**). Interestingly, the function of type I interferon has been well documented to be essential for controlling viral infection (PERRY et al., 2005), and may therefore be reflective of a stronger infection in patients further along the M1 axis, which is confirmed by the simultaneously increasing disease severity.

This type of cross-data-type integrated analysis can suggest experimentally testable hypotheses that might lead to improved treatment. For example, is there a level of IDO inhibition that, if provided to the right patients at the right time, might help restore a more balanced and functional anti-SARS-CoV-2 immune response while reducing hypotension? This comprehensive data set can be explored for testing or generating a number of other hypotheses, including detailed analyses of immune responses to investigational antiviral or immunomodulatory therapies for COVID-19. However, a limitation is that the 50-sample, 26-patient cohort studied here is relatively modest, the numbers of co-morbidities present in this population is large, and the time-trajectories that are studied are relatively short, and so don’t capture the resolution of the infection. Nevertheless, our dataset establishes an unprecedented level of detail on the impacts of COVID-19 on the immune systems. We show that the collected single cell, bulk plasma, and clinical data sets can be synergized to reveal clear interpretable biological trends that can be associated with COVID-19 patient outcomes and may be suggestive of treatment strategies. This broad systems immunology approach should be applicable towards understanding immune responses in a host of other infectious diseases.

## ACKNOWLEGEMENT

We are grateful to all participants and blood donors in this study. We are extremely grateful to the medical teams at Swedish Medical Center for their support of this research and for their assuming personal risk in their tireless care for patients suffering from COVID-19. We thank the Northwest Genomic Center, especially, Debbie Nickerson, Erica Ryke, and Peter Anderson for the help with sequencing services. We thank the insightful discussion from Prof. David Baltimore, Prof. Ilya Shmulevich and the ISB COVID19 Study Group: Inyoul Lee, Kay Chinn, Scott Bloom, Guenther Kahlert, David Baxter, Zac Simon, Matt Idso, Rachel Calder, William Chour, Alphonsus Ng, Jimmi Hopkins, John Heath, Jingyi Xie, John Heath, Jessica Yee, Kari Tetrault, Lee Rowen, Rachel Liu, Rongyu Zhang, Shannon Fallen, Simran Sidhu, Yue Lu, David Gibbs, Michael Strasser; and the Swedish COVID-19 Research Steering Committee: John Pauk, Bill Berrington, Michael Bolton, Mary Micikas, Sonam Nyatsatsang, Cynthia Maree, Shane O’Mahony, Kelly Sweerus, Anne Lipke, George Pappas, Mark Sullivan, Karen Koo, Chris Dale, George Lopez, Naomi Diggs, Hank Kaplan, Krish Patel, Livia Hegerova, Vanessa Dunleavy, Phil Gold, John Pagel, Joshua Mark, Doug Kieper, Jim Scanlan, Evonne Lackey, Jodie Davila, Justin Rueda, Julie Wallick, Heather Algren, Jennifer Hansberry and Elizabeth Wako.

The ISB-Swedish COVID19 Biobanking Unit: Rick Edmark, John Heath, Simran Sidu, Jessica Yee, Cara McCoy, Jeremy Johnson, Renee Duprel, Audri Hubbard, Theresa Davis, Julie Thatcher and Zuraya Aziz, Thea Swanson, Yong Zhou, Lesley Jones, Sarah Li, Paula Manner, Andrea Drouhard, Stephanie Johnson, Julia Karr, Clementine Chalal, Mattie Sader, Margo Badman, Allison Everett, Adel Islam, Jodie Davila and Julie Wallick.

We gratefully acknowledge funding support from the Wilke Family Foundation (J.R.H.), the Murdock Trust (J.R.H.), the Swedish Medical Center Foundation (J.D.G.), the Parker Institute for Cancer Immunotherapy (J.R.H., M.M.D., P.G., L.L.L., J.A.B.), Merck and the Biomedical Advanced Research and Development Authority under Contract HHSO10201600031C (J.R.H.). K.W. was funded by DOD W911NF-17-2-0086, NIH R01 DA040395, and NIH UG3TR002884. J.H. was funded by National Center For Advancing Translational Sciences of the National Institutes of Health under Award Number OT2 TR003443. R.G. was funded by the NIH Human Immunology Project Consortium (U19AI128914) and the Vaccine and Immunology Statistical Center (Bill and Melinda Gates Foundation grant no. OPP1032317). Further funding by NIH AI068129 (L.L.L.), and NIH R21 AI138258 (N.S.).

## AUTHOR CONTRIBUTIONS

Conceptualization, Y.S., J.D.G.., and J.R.H.; Resources, J.D.G., and J.R.H.; Methodology, Y.S., D.C., J.S., and J.R.H.; Investigation Y.S., D.C., C.L., D.Y., J.C., C.D., V.V., K.S., P.T., V.D., P. B., G.Q., B.S., S.K., C.R., A.X., J.L., S.D., A.R., J.Z., K.M., R.E., S.H., L.J., Y.Z., R.R., S.M., S.M., C.D., J.W., H.A., Z.M., A.M., W.W., N.D.P., S.H., N.S., K.W., J.H., L.H., A.A., J.A.B., L.L.L, P.G., R.G., M.M.D., J.D.G., and J.R.H.; Formal Analysis, Y. S., D.C., D.Y., C.L., C.D., V.V., V.D., R.G., and J.R.H.; Writing– Original Draft, Y.S., D.C., D.Y., and J.R.H.; Writing – Review & Editing, Y.S., D.C., C.L., D.Y., J.C., C.D., V.V., K.S., P.T., V.D., P. B., G.Q., B.S., S.K., C.R., A.X., A.R., J.Z., A.M., W.W., N.D.P., S.H., K.W., N.S., J.H., L.H., A.A., J.A.B., L.L.L, P.G., R.G., M.M.D., J.D.G., and J.R.H.

## SI Figures

Figure S1. Overview of the multi-omic characterization of immune responses in COVID-19 patients.

A. The swimmers’ plot of WHO severity scores for study patients. The scores were derived from electronic health records. Symptom onset and pre-hospitalization are indicated by black rectangles and lines. WHO scores are presented at 6-hour intervals during hospitalization. Blood draws are indicated by upside-down blue triangles and medication administration by symbols overlaid on the colored bands.

B. Box plots of clinical data comparing non-ICU (yellow, n=34) and ICU (red, n=16) patient values. Ranges that specify normal limits are indicated by the blue (lower) and orange (upper) dashed lines.

C. Correlation matrix of 20 clinical features and 8 blood cell-type counts from 50 COVID-19 patients. The square size corresponds to absolute value of correlation coefficient, with blue (red) color indicating a positive (negative) correlation. Significance is indicated by: (*p < 0.05, **p< 0.01, ***p < 0.001).

D. Spearman correlations between the WHO_ordinal_scale of disease severity with principal component (PC) 1 of plasma proteomic (left) and PC1 of plasma metabolomic (right) data. The regression line is drawn with the 95% confidence area shaded. Spearman Correlation coefficient and associated p-value shown.

E. Box plot of plasma protein concentrations in healthy donors (green), non-ICU blood draws (yellow), and ICU blood draws (red) of COVID-19 patients.

F. Heatmap showing gene expression of well-known markers specific for each cell type. Clear separation of annotated cell type is observed. The color of the column and row markers corresponds to the cluster colors of the UMAP shown in Fig 1E.

G. Heatmap showing levels of well-known surface proteins specific for each cell type. Clear separation of annotated cell type is observed. The color of the column and row markers corresponds to the cluster colors of the UMAP shown in Fig 1E.

H. A bar plot showing the relative proportion of major immune cell type across each sample, as determined from the analysis of the 10x single cell transcriptome and proteomic data set.

I. Box plot depicting the proportions selected immune cell types within PBMCs from healthy donors (green), non-ICU blood draws (yellow), and ICU blood draws (red) of COVID-19 patients.

Figure S2. CD8+ T cell heterogeneity in COVID-19 patients and its association with disease severity.

A. UMAP embedding of all CD8+ T cells colored by unsupervised clustering (top left panel) and expression of selected genes (other panels).

B. UMAP embedding of CD8+ T cells colored by the levels of selected surface proteins, annotated with the cell state that they represent.

C. UMAP embedding of all CD8+ T cells colored by the density of cells characterized by different clonal expansion sizes: n=1, n=2-4, and n>=5.

D. Clonal expansion status of each CD8+ T cell subset from unsupervised clustering. Bar plot shows the normalized clonal composition.

E. Boxplots showing the mRNA expression levels of 3 transcripts in healthy donors (green), non-ICU blood draws (yellow), and ICU blood draws (red) of COVID-19 patients.

F. Scatter plots showing the naïve gene signature score (x-axis) and cytotoxic gene signature score (y-axis) of individual CD8+ T cells from all PBMC samples. Plots colored according to unsupervised clustering (from Figure 2A), patient status from whom individual cells were drawn, proliferation gene signature score, and exhaustion gene signature score are color-coded in each panel.

Figure S3. CD4+ T cells display heterogeneous proliferation, cytotoxic activity and clonal expansion levels and are associated with the severity of COVID-19.

A. UMAP embedding of all CD4+ T cells colored by unsupervised clustering (top left panel) and expression of other representative genes (other panels).

B. UMAP embedding of CD4+ T cells shaded by the levels of selected surface proteins.

C. UMAP embedding of all CD8+ T cells colored by the density of cells characterized by different clonal expansion sizes: n=1, n=2-4, and n>=5.

D. Clonal expansion status of each CD4+ T cell subset from unsupervised clustering. Bar plot shows the normalized clonal composition.

E. Boxplots showing the mRNA expression levels of three representative transcripts from CD8+ T cells in healthy donors (green), non-ICU blood draws (yellow) and ICU blood draws (red) of COVID-19 patients.

F. Functional characterization of CD4+ T cells from PBMC samples using single-cell secretome analysis. Single-cell polyfunctional strength index (PSI) of CD4+ T cells from healthy donors, non-ICU blood draws (yellow), and ICU blood draws (red) of COVID-19 patients. The PSI is computed for five categories of cytokines separately, as indicated by the box plot title.

Figure S4. Deep profiling of COVID-19 patient B cell populations reveals heterogeneous differentiation status that correlates with disease severity.

A. UMAP embedding of all B cells colored by unsupervised clustering (top left panel) and other representative genes (other panels).

B. UMAP embedding of all B cells colored by levels of one representative surface protein.

C-E. Boxplots showing the, mRNA expression levels (panel C), surface protein levels (panel D), and the pathway enrichment scores (panel E) in healthy donors (green), non-ICU blood draws (yellow), and ICU blood draws (red) of COVID-19 patients.

Figure S5. Alterations of antigen presentation and inflammation in monocytes are associated with COVID-19 disease severity.

A. UMAP embedding of all monocytes colored by unsupervised clustering (top left panel) and other representative genes (other panels).

B. UMAP embedding of monocytes colored by representative pathway enrichment scores.

C. Functional characterization of monocytes from PBMC samples using single-cell secretome analysis. Boxplots indicate the percentage of monocytes secreting IL6, IL8, MCP1, MIP1-A, MIP1-B, and TNF-α from samples in healthy donors (green), non-ICU blood draws (yellow), and ICU blood draws (red) of COVID-19 patients.

D. Spearman correlation of monocytes MHC class II gene expression with plasma IL6 level. Regression line indicated in black, with a 95% confidence interval shown in shaded gray. Spearman Correlation coefficient and associated p-value shown.

Figure S6. Deep profiling of COVID-19 patient NK cell populations reveals heterogeneous proliferation that correlates with disease severity.

A. UMAP embedding of all NK cells colored by unsupervised clustering (top left panel) and other representative genes (other panels).

B. UMAP embedding of all NK cells colored by levels of two representative surface proteins.

C. UMAP embedding of all NK cells colored by embedding density of cells from different groups of blood draw samples (healthy, non-ICU, and ICU). Two clusters that displayed significant changes of density from group to group are circled using dashed lines color-coded for the cluster color of panel A.

D. Box plots depicting the kinetic changes (T2-T1) of the levels of mRNA in NK cells between the 1st and 2nd blood draws from patients who improved (green) and patients who did not (red).

Figure S7. Multi-omic data integrating across cell types resolve an orchestrated response of innate and adaptive immune cell gene module that correlated with disease severity.

A. Spearman correlations of gene module (M)1 score with a few representative plasma cytokines. The regression line is indicated in black, with the 95% confidence area shown in shaded gray. Spearman Correlation coefficient and associated p-value shown.

B. Spearman correlations of M1 score with a few representative plasma metabolites. The regression line is indicated in black, with the 95% confidence area shown in shaded gray. Spearman Correlation coefficient and associated p-value shown.

C. Correlation matrix of 21 clinical labs and M1 score in 50 COVID-19 patient samples. The square size corresponds to absolute value of the correlation coefficient, and the color corresponds to correlation value with blue (red) indicating positive (negative) correlation. Significance indicated by (*p<0.05, **p<0.01, ***p <0.001).

D. Spearman correlations of the M1 score with levels of a few representative mRNA, surface proteins and pathway enrichment scores across different immune cell types. The regression line is indicated in cell-type specific color, with the 95% confidence area shown in shaded cell-type specific color. Spearman Correlation coefficient and associated p-value shown.

## METHOD DETAILS

### Quantify disease severity with World Health Organization (WHO) Ordinal Scale

Severity of COVID-19 was clinical status assessed throughout a patient’s encounter in accordance to the 9-point WHO Ordinal Scale for Clinical Improvement consisting of the following categories: 0) uninfected - no evidence of infection; 1) ambulatory - no limitation of activities; 2) ambulatory - limitation of activities; 3) hospitalized, mild - no oxygen therapy; 4) hospitalized, mild - oxygen by mask or nasal prongs; 5) hospitalized, severe - non-invasive ventilation or high-flow oxygen; 6) hospitalized, severe - intubation and mechanical ventilation; 7) hospitalized, severe - ventilation + additional organ support; and 8) dead – death (WHO, 2020).

### EHR clinical data analysis and designation of critical illness status

WHO grades for time of blood draw were determined by manual expert review. WHO grades for Figure S1A were automatically generated from data extracted from the electronic health record, and plotted for 6-hour time intervals based on end-interval grade. Automated results were compared against manual expert review for 50% of study subjects.

The following data were collected from the subject’s electronic health record (EHR): complete blood count (CBC) with differential, comprehensive metabolic panel. Lab data were extracted from the nearest time point to the each blood draw, if available within a window +/- two days. First blood draw (n=26), second blood draw (n=24). Estimated absolute numbers of B cell, CD4^+^ T cell, CD8^+^ T cells and NK cells, were derived using observed percentages of lymphocytes from study blood draws, multiplied by the total lymphocyte count from the EHR labs described above. Blood draws were classified as non-ICU (n=34) and ICU (n=16). The physical location and the ICU admission status were concordant at the time of each blood draw **(Table S1A)**. We used unpaired t-test to determine the statistical significance between ICU and non-ICU patients.

### Correlation between module1 score (M1) with EHR labs and CBCs

We conducted Pearson correlation analysis to evaluate the association between gene module 1 and labs as well as CBCs.

### Plasma and PBMC Isolation

Plasma and PBMC isolation was conducted with standard protocols from Bloodworks Northwest (Seattle, WA). Patient blood were collected in BD Vacutainer (EDTA) tubes (Becton, Dickinson and Company, Franklin Lakes, NJ). Plasma fractions were collected after centrifuged at 800 x g at 4°C for 10 min, aliquoted and stored until use at −80°C. The rest of the blood were diluted with PBS (pH7.2) to 2X of original volume and layered over 15 ml Ficoll (GE Healthcare, Waukesha, WI) in SepMate-50 tubes (Vancouver, BC). After centrifuged at 800 x g for 15 min at room temperature, the PBMC layer was poured into a 50 ml conical tube. The cells were washed twice with autoMACS Rinsing Solution (Miltenyi Biotec, Auburn, CA) and centrifuge at 250 x g for 10 min, at RT. PBMC pellets were gently resuspend in 5 ml Rinsing Solution and a 5 μl aliquot was diluted 1:10 v/v for cell counting. Cells in 18 μl of diluted samples were first mixed with 2 μl of Acridine Orange / Propidium Iodide Stain (Logos Biosystems, Annandale, VA), 10 μl was then loaded to a PhotonSlide (Logos Biosystems) and counted in a LUNA FL Dual Fluorescence cell counter (Logos Bioystems). Cryopreservation freeze media CryoStor CS-10 (Biolife Solutions, Bothell, WA) was slowly added to make a concentration of 2.5 million PBMC/ml. Cells were aliquoted in Cryotube vials (ThermoFisher, Waltham, MA) and frozen in CoolCell LX Cell Freezing Container (Corning, Corning, NY) at −80°C for at least 2 hours before stored until use in LN.

### Plasma proteomics

Plasma concentrations of proteins were measured using the ProSeek Cardiovascular II, Inflammation, Metabolism, Immune Response, and Organ Damage panels (Olink Biosciences, Uppsala, Sweden). Health control plasma samples were processed at Olink facilities in Boston, MA. Plasma samples from COVID-19 participants were assayed at the Institute for Systems Biology. Proteins from patient plasma were measured using proximity extension assay (PEA) (Olink Proteomics, Uppsala, Sweden) which allows for the simultaneous analysis of 92 protein biomarkers on each panel. Five panels including Inflammation, Cardiovascular II, Organ Damage, Immune Response and Metabolism were run using 82 patient plasma samples as well as 8 replicates of a pooled healthy control. One microliter of plasma was incubated overnight and allowed to bind with oligonucleotide-labeled antibody pairs to form specific DNA duplexes. This template was then extended and pre-amplified, and the individual protein markers were measured using high-throughput microfluidic real-time PCR. The resulting Ct values were normalized against an extension control, an inter-plate control, and adjusted with a correction factor according to the manufacturer’s instructions to calculate a normalized protein expression value (NPX) in log2 scale.

Samples were processed in batches with pooled quality control samples included in each batch; potential batch effects were subsequently adjusted using the pooled control samples, as previously described (Manor et al., 2018). For analysis, a threshold of less than 25% missing values was set for each protein. Missing values for the proteins were imputed to be the limits of detection. A total of 454 proteins were included in further analyses.

### Plasma metabolomics

Metabolon (Morrisville, NC, USA) conducted the metabolomics assays for all participant plasma samples used in this study. Data were generated with the Global Metabolomics platform via ultra-high-performance liquid chromatography/tandem accurate mass spectrometry. Sample handling, quality control, and data extraction, along with biochemical identification, data curation, quantification, and data normalizations have been previously described (Manor et al., 2018; Wittmann et al., 2014). Samples were processed in batches with pooled quality control samples included in each batch; pooled quality control samples were consistent across batches. Potential batch effects for each metabolite were adjusted by dividing by the corresponding average value identified in the pooled quality control samples from the same batch. For analysis, the raw metabolomics data were median scaled within each batch such that the median value for each metabolite was one. Metabolite levels were further quality controlled by removing metabolites that were observed in less than 75% of samples. Missing values for remaining metabolites were then imputed using the minimum observed value for each metabolite. Values for each metabolite were subsequently log transformed. 847 different plasma metabolites were used for downstream statistical analyses.

### Principal component analysis (PCA) of plasma proteomic and metabolomics data

PCA was performed separately for metabolomics and proteomic data using all metabolites or proteins that passed quality control, respectively. Values were centered and scaled prior to dimensional reduction. PCA plots were colored by the severity of the patient based on the WHO Ordinal Scale at the time of the blood draw.

### Statistical analysis of plasma proteomic and metabolomics data

#### Association with COVID-19 status

We assessed differential abundance of protein and metabolite levels between healthy and all COVID-19 participants. Levels of proteins and metabolites were first adjusted for age and sex. Two-sided Mann-Whitney U tests were performed between all health control samples and all COVID-19 samples for proteins and metabolites separately. Multiple testing correction was subsequently applied using the Benjamini-Hochberg (BH) procedure with false discovery rate (FDR) at 5%. We also performed univariate logistic regression between COVID-19 status and adjusted levels of metabolites and protein levels separately to calculate the log odds ratio.

#### Association with COVID-19 severity

Spearman rank correlation was performed to assess the association of metabolite and protein levels with the WHO Ordinal Scale score of the patient at the time of the sample. Age and sex adjusted protein and metabolite levels were tested separately and multiple testing correction was performed using BH FDR <5%. We also assessed associations between adjusted protein and metabolite levels with the participant location at the time of blood draw—intensive care unit (ICU), non-ICU hospitalized, and healthy/uninfected—using two-sided Mann-Whitney U tests. Similarly, multiple testing correction using BH FDR <5% was applied.

#### Association with gene module1 score of single cell data

Spearman rank correlation was also performed to test the association of adjusted metabolite and protein levels with the M1 score derived from surprisal analysis of single cell data. BH FDR <5% was subsequently used.

### Single-cell multiplex secretome assay of immune cells

Cryopreserved PBMCs were thawed and incubated in the RPMI 1640 for overnight recovery at 37°C, 5% CO2. Recovered cell viability was measured at >95% for all samples. After overnight recovery, CD4+ and CD8^+^ T cell populations and monocytes were magnetically isolated using the CD4+, CD8+ Microbeads and Pan Monocyte Isolation Kit (Miltenyi Biotec, Bergisch Gladbach, Germany) sequentially.

The enriched CD4^+^ and CD8^+^ T cells (100,000 cells/well in a 96 well-plate) were stimulated for 6 hours with plate-bound anti-CD3 antibodies (pre-coated at 10 μg/mL overnight at 4°C) and soluble anti-CD28 antibodies (5 μg/mL) in complete RPMI-1640 at 37 °C, 5% CO2. The enriched monocytes at 1 x 105/ml were stimulated with 10 ng/ml LPS for 12 hours.

After stimulation, the activated cells were collected, washed, and stained with membrane stain (included in the IsoPlexis kit). The stained cells were then loaded onto the chip consisting of the 12,000 chambers pre-coated with an array of 32 cytokine capture antibodies. The chip was inserted into the fully automated IsoLight for further incubation for 16 hours and the cytokines were detected by a cocktail of detection antibodies followed by the fluorescent labeling. The scanned fluorescent signal was analyzed by IsoSpeak software to calculate the numbers of cytokine-secreting cells, the intensity level of cytokines and polyfunctional strength index (PSI).

See the below for the measured cytokines in each panel.

Single-Cell Adaptive Immune cytokine panel including the following subsets of cytokines. Effector: Granzyme B, IFN-γ, MIP-1α, Perforin, TNF-α, TNF-β; Stimulatory: GM-CSF, IL-2, IL-5, IL-7, IL-8, IL-9, IL-12, IL-15, IL-21; Chemoattractive: CCL11, IP-10, MIP-1β, RANTES; Regulatory: IL-4, IL-10, IL-13, IL-22, TGFβ1, sCD137, sCD40L; Inflammatory: IL-1β, IL-6, IL-17A, IL-17F, MCP-1, MCP-4.

Single-Cell Innate Immune cytokine panel including the following subsets of cytokines. Effector: IFN-γ, MIP-1α, TNF-α, TNF-β; Stimulatory: GM-CSF, IL-8, IL-9, IL-15, IL-18, TGF-α, IL-5; Chemoattractive: CCL11, IP-10, MIP-1β, RANTES, BCA-1; Regulatory: IL-10, IL-13, IL-22, sCD40L; Inflammatory: IL-1β, IL-6, IL-12-p40, IL-12, IL-17A, IL-17F, MCP-1, MCP-4, MIF; Growth Factors: EGF, PDGF-BB, VEGF.

### Single cell multi-omics assay

Chromium Single Cell Kits (10x Genomics) were utilized to analyze the transcriptomic, surface protein levels and TCR and BCR sequences simultaneously from the same cell. Experiments were performed according to manufacturer’s instructions.

Briefly, cryopreserved PBMCs were thawed and 1X red blood cell lysis solution (Biolegend) was used to lyse any remaining red blood cells in the PBMC samples. Cells were stained with custom TotalSeq-C human antibodies (BioLegend) before loading onto a Chromium Next GEM chip G. Cells were lysed for reverse transcription and complementary DNA (cDNA) amplification in the Chromium Controller (10X Genomics). The polyadenylated transcripts were reverse-transcribed inside each gel bead-in-emulsion afterwards. Full-length cDNA along with cell barcode identifiers were PCR-amplified and sequencing libraries were prepared and normalized. The constructed library was sequenced on Novaseq platform (Illumina).

### Single cell RNA-seq data processing

Droplet-based sequencing data were aligned and quantified using the Cell Ranger Single-Cell Software Suite (version 3.0.0, 10x Genomics) against the GRCh38 human reference genome. Quality of cells in each demultiplexed sample were then assessed based on three metrics: (1)The number of total Unique molecular identifiers (UMI) counts per cell (library size); (2)the number of detected genes per cell; and (3)the proportion of mitochondrial gene counts. Doublets were simultaneously identified in sample demultiplexing and, were filtered out prior to filtering based on the aforementioned three metrics. After QC metric filtering, a total of 221,748 cells were retained for downstream analysis. Scanpy (Wolf et al., 2018) was used to normalize cells via CPM normalization (UMI total count of each cell was set to 10^6^) and log1p transformation (natural log of CPM plus one).

### Single cell RNA-seq dimension reduction and unsupervised clustering

PCA was performed on the gene expression matrix and Scanpy was used to build a nearest neighbor graph to calculate a UMAP and find clusters via the louvain algorithm. Clusters identified in this first round of clustering were annotated based on the expression of canonical marker genes. Clusters that were not uniform in their expression of well-known marker genes were extracted and a second round of dimension reduction and clustering was performed on these subsets, this was mainly to separate T cells and NK cells. The surface protein level matrix of identified T cells from the first and second round of clustering were subject to the same dimension reduction and a CD4 surface protein x CD8 surface protein scatterplot was utilized to linearly separate the T cells into CD4+ T cells and CD8+ T cells, T cells that had zero expression of both CD4 and CD8 surface proteins were categorized as Other T cells. Major immune cell types CD4+ T cells, CD8+ T cells, monocytes, B cells, and NK cells were further analyzed via a construction of their single-cell regulatory networks by inputting their normalized gene expression matrices into pySCENIC (Van de Sande et al., 2020). The resultant single-cell transcription factor modules were utilized for cell type specific dimension reduction and clustering via either the louvain (CD8+ T cells) or leiden algorithm (all other major cell types). All reduced dimensions and clusters were calculated by Scanpy. Surprisal analysis was applied as described previously (Remacle et al., 2010; Zadran et al., 2014). Briefly, for each PBMC sample n, the measured, from certain cell type, average expression level of mRNA i, ln Xi(n), was expressed as a sum of a steady state term and the major constraints λj(n)Gij representing deviations from the steady state. Each deviation term was a product of a sample-dependent weight of the constraint λj(n), and the sample-independent contribution of the transcript to that constraint Gij. In this study, the most dominant constraint, λ1(n) was investigated as the immune response module M1.

### Single-cell multi-omics disease severity analysis

The data set collected here can be analyzed, at a high level, in two different ways. First, each blood draw corresponds to a time point at which that patient was characterized by a level of disease severity. This can be approximated by whether the patient was in the intensive care unit (ICU) or not (**Table S1**) or, at higher resolution, by using the World Health Organization (WHO) Ordinal Scale of 0-8 for COVID-19, see above (**Table S1**). WHO score was calculated in 6 hour intervals, and the score was used that corresponded to the window in which the blood was drawn. Grouping each blood draw according to a contemporary measure of disease severity can provide insight into how various immune cell populations, regulatory networks, and other biomarkers vary with disease severity level by sample. A second view is to consider each patient separately, so that the two blood draws are used to capture an individual patient’s disease trajectory. Here, patients that improve between T1 and T2 can be differentiated from patients advancing to more severe infection. We utilize both types of analyses here. Disease trajectories, including time points of therapeutic interventions, the time of the T1 and T2 blood draws, and WHO ordinal scale measures of disease severity, are provided in Figure S1A. These trajectories start at the self-reported day of symptom onset and give WHO score during time of hospital admission.

### Statistical analysis of single-cell multi-omics data

Differential expression analysis was performed using the R package MAST (Finak et al., 2015). For each cell type (CD8^+^ T cells, CD4^+^Tcells, monocytes, B cells, and NK cells), we investigated the interaction model between the patient location (ICU or non-ICU) and time with T1 and non-ICU as baselines in samples not treated with steroid or tocilizumab. The normalized gene-cell barcode was used as input. These models included as covariates the batch and patient ID, as well as the cellular detection rate (CDR) to correct for biological and technical nuisance factors. Multiple hypothesis testing correction was performed by controlling the false discovery rate (FDR). Genes were declared significantly differentially expressed at a FDR of 5% and a fold-change > 1.5.

Gene set enrichment analysis (GSEA) was performed using the R package MAST with the bootVcov1 R function. The same models as above were investigated. A FDR cutoff of 5% was used to determine significant gene sets. KEGG, Hallmark, and blood transcription modules (Li et al., 2014) were used as gene sets. KEGG and Hallmark gene sets were downloaded from MSigDB database (Liberzon et al., 2011).

### Gene set variation analysis (GSVA)

To quantify the pathway enrichment score across samples or cells, GSVA was performed using the R package GSVA (v.3.11) (Hänzelmann et al., 2013) to identify the most changed pathways between samples/cells with the averaged log_2_ (TPM+1) data as input.

### Single cell TCR-seq data processing and RNA-seq integration

Droplet-based sequencing data were aligned and quantified using the Cell Ranger Single-Cell Software Suite (version 3.1.0, 10x Genomics) against the GRCh38 human VDJ reference genome. Resultant filtered annotated contigs were analyzed via Scirpy (Sturm et al., 2020). The aforementioned files were read in as TCRs, filtered for either CD4^+^ or CD8^+^ T cells, as identified via single cell RNA-seq analysis, and then subject to clonotype definition and clonal expansion analysis. Samples were then concatenated together and merged with gene expression data for simultaneous single cell TCR and RNA data visualization.

## Lead Contact

Further information and requests for resources and reagents should be directed to and will be fulfilled by the Lead Contact, Dr. James R. Heath.

## Data Availability

Due to potential risk of de-identification of pseudonymized RNA sequencing data the raw data will be available under controlled access in the EGA repository upon journal acceptance. In addition, processed data are available at Array Express, accession number: E-MTAB-9357. For access to data prior to journal acceptance, please contact the corresponding authors.

All data will be integrated for further visualization and analysis in ISB COVID-19 data explorer at https://atlas.fredhutch.org/isb/covid/.

## REFERENCES

Abel, B., Tameris, M., Mansoor, N., Gelderbloem, S., Hughes, J., Abrahams, D., Makhethe, L., Erasmus, M., de Kock, M., van der Merwe, L., et al. (2010). The novel tuberculosis vaccine, AERAS-402, induces robust and polyfunctional CD4+ and CD8+ T cells in afile:///Users/danyuan/Downloads/26046523.nbibdults. Am. J. Respir. Crit. Care Med. 181, 1407–1417.

Agresta, L., Hoebe, K.H.N., and Janssen, E.M. (2018). The Emerging Role of CD244 Signaling in Immune Cells of the Tumor Microenvironment. Front. Immunol. 9, 2809.

Becht, E., McInnes, L., Healy, J., Dutertre, C.-A., Kwok, I.W.H., Ng, L.G., Ginhoux, F., and Newell, E.W. (2019). Dimensionality reduction for visualizing single-cell data using UMAP. Nat. Biotechnol. 37, 38–44.

Belladonna, M.L., Orabona, C., Grohmann, U., and Puccetti, P. (2009). TGF-β and kynurenines as the key to infectious tolerance. Trends Mol. Med. 15, 41–49.

Bhatraju, P.K., Ghassemieh, B.J., Nichols, M., Kim, R., Jerome, K.R., Nalla, A.K., Greninger, A.L., Pipavath, S., Wurfel, M.M., Evans, L., et al. (2020). Covid-19 in Critically Ill Patients in the Seattle Region — Case Series. N. Engl. J. Med. 382, 2012–2022.

Boyd, A., Almeida, J.R., Darrah, P.A., Sauce, D., Seder, R.A., Appay, V., Gorochov, G., and Larsen, M. (2015). Pathogen-Specific T Cell Polyfunctionality Is a Correlate of T Cell Efficacy and Immune Protection. PLoS One 10, e0128714.

Cao, X. (2020). COVID-19: immunopathology and its implications for therapy. Nat. Rev. Immunol. 20, 269–270.

Chen, Y., Feng, Z., Diao, B., Wang, R., Wang, G., Wang, C., Tan, Y., Liu, L., Wang, C., Liu, Y., et al. (2020). The Novel Severe Acute Respiratory Syndrome Coronavirus 2 (SARS-CoV-2) Directly Decimates Human Spleens and Lymph Nodes. MedRxiv 2020.03.27.20045427.

Chua, R.L., Lukassen, S., Trump, S., Hennig, B.P., Wendisch, D., Pott, F., Debnath, O., Thürmann, L., Kurth, F., Völker, M.T., et al. (2020). COVID-19 severity correlates with airway epithelium–immune cell interactions identified by single-cell analysis. Nat. Biotechnol.

Cicko, S., Grimm, M., Ayata, K., Beckert, J., Meyer, A., Hossfeld, M., Zissel, G., Idzko, M., and Müller, T. (2015). Uridine supplementation exerts anti-inflammatory and anti-fibrotic effects in an animal model of pulmonary fibrosis. Respir. Res. 16, 105.

Diao, B., Wang, C., Tan, Y., Chen, X., Liu, Y., Ning, L., Chen, L., Li, M., Liu, Y., Wang, G., et al. (2020). Reduction and Functional Exhaustion of T Cells in Patients With Coronavirus Disease 2019 (COVID-19). Front. Immunol. 11, 827.

Finak, G., McDavid, A., Yajima, M., Deng, J., Gersuk, V., Shalek, A.K., Slichter, C.K., Miller, H.W., McElrath, M.J., Prlic, M., et al. (2015). MAST: a flexible statistical framework for assessing transcriptional changes and characterizing heterogeneity in single-cell RNA sequencing data. Genome Biol. 16, 278.

Giamarellos-Bourboulis, E.J., Netea, M.G., Rovina, N., Akinosoglou, K., Antoniadou, A., Antonakos, N., Damoraki, G., Gkavogianni, T., Adami, M.-E., Katsaounou, P., et al. (2020). Complex Immune Dysregulation in COVID-19 Patients with Severe Respiratory Failure. Cell Host Microbe 27, 992–1000.e3.

Hansen, A.M., Driussi, C., Turner, V., Takikawa, O., and Hunt, N.H. (2000). Tissue distribution of indoleamine 2,3-dioxygenase in normal and malaria-infected tissue. Redox Rep. 5, 112–115.

Hänzelmann, S., Castelo, R., and Guinney, J. (2013). GSVA: gene set variation analysis for microarray and RNA-Seq data. BMC Bioinformatics 14, 7.

Herold, T., Jurinovic, V., Arnreich, C., Lipworth, B.J., Hellmuth, J.C., Bergwelt-Baildon, M. von, Klein, M., and Weinberger, T. (2020). Elevated levels of IL-6 and CRP predict the need for mechanical ventilation in COVID-19. J. Allergy Clin. Immunol. 146, 128–136.e4.

Ivashkiv, L.B. (2020). The hypoxia–lactate axis tempers inflammation. Nat. Rev. Immunol. 20, 85–86.

Iype, E., and Gulati, S. (2020). Understanding the asymmetric spread and case fatality rate (CFR) for COVID-19 among countries. MedRxiv.

St. John, A.L., and Rathore, A.P.S. (2019). Adaptive immune responses to primary and secondary dengue virus infections. Nat. Rev. Immunol. 19, 218–230.

Juno, J.A., van Bockel, D., Kent, S.J., Kelleher, A.D., Zaunders, J.J., and Munier, C.M.L. (2017). Cytotoxic CD4 T Cells-Friend or Foe during Viral Infection? Front. Immunol. 8, 19.

Kravchenko-Balasha, N., Wang, J., Remacle, F., Levine, R.D., and Heath, J.R. (2014). Glioblastoma cellular architectures are predicted through the characterization of two-cell interactions. Proc. Natl. Acad. Sci. U. S. A. 111, 6521–6526.

Kravchenko-Balasha, N., Shin, Y.S., Sutherland, A., Levine, R.D., and Heath, J.R. (2016). Intercellular signaling through secreted proteins induces free-energy gradient-directed cell movement. Proc. Natl. Acad. Sci. U. S. A. 113, 5520–5525.

Li, S., Rouphael, N., Duraisingham, S., Romero-Steiner, S., Presnell, S., Davis, C., Schmidt, D.S., Johnson, S.E., Milton, A., Rajam, G., et al. (2014). Molecular signatures of antibody responses derived from a systems biology study of five human vaccines. Nat. Immunol. 15, 195–204.

Liberzon, A., Subramanian, A., Pinchback, R., Thorvaldsdóttir, H., Tamayo, P., and Mesirov, J.P. (2011). Molecular signatures database (MSigDB) 3.0. Bioinformatics 27, 1739–1740.

Liu, L., Du, X., Zhang, Z., and Zhou, J. (2018). Trigonelline inhibits caspase 3 to protect β cells apoptosis in streptozotocin-induced type 1 diabetic mice. Eur. J. Pharmacol. 836, 115–121.

Lu, Y., Xue, Q., Eisele, M.R., Sulistijo, E.S., Brower, K., Han, L., Amir, E.D., Pe\textquoterighter, D., Miller-Jensen, K., and Fan, R. (2015). Highly multiplexed profiling of single-cell effector functions reveals deep functional heterogeneity in response to pathogenic ligands. Proc. Natl. Acad. Sci. 112, E607–E615.

Lukaszewicz, A.-C., Grienay, M., Resche-Rigon, M., Pirracchio, R., Faivre, V., Boval, B., and Payen, D. (2009). Monocytic HLA-DR expression in intensive care patients: Interest for prognosis and secondary infection prediction *. Crit. Care Med. 37.

Ma, C., Fan, R., Ahmad, H., Shi, Q., Comin-Anduix, B., Chodon, T., Koya, R.C., Liu, C.-C., Kwong, G.A., Radu, C.G., et al. (2011). A clinical microchip for evaluation of single immune cells reveals high functional heterogeneity in phenotypically similar T cells. Nat. Med. 17, 738–743.

Ma, C., Cheung, A.F., Chodon, T., Koya, R.C., Wu, Z., Ng, C., Avramis, E., Cochran, A.J., Witte, O.N., Baltimore, D., et al. (2013). Multifunctional T-cell analyses to study response and progression in adoptive cell transfer immunotherapy. Cancer Discov. 3, 418–429.

Manor, O., Zubair, N., Conomos, M.P., Xu, X., Rohwer, J.E., Krafft, C.E., Lovejoy, J.C., and Magis, A.T. (2018). A Multi-omic Association Study of Trimethylamine N-Oxide. Cell Rep. 24, 935–946.

Marques, E.P., Ferreira, F.S., Santos, T.M., Prezzi, C.A., Martins, L.A.M., Bobermin, L.D., Quincozes-Santos, A., and Wyse, A.T.S. (2019). Cross-talk between guanidinoacetate neurotoxicity, memory and possible neuroprotective role of creatine. Biochim. Biophys. Acta. Mol. Basis Dis. 1865, 165529.

Mathew, D., Giles, J.R., Baxter, A.E., Greenplate, A.R., Wu, J.E., Alanio, C., Oldridge, D.A., Kuri-Cervantes, L., Pampena, M.B., D’Andrea, K., et al. (2020). Deep immune profiling of COVID-19 patients reveals patient heterogeneity and distinct immunotypes with implications for therapeutic interventions. BioRxiv Prepr. Serv. Biol. 2020.05.20.106401.

Moon, C. (2020). Fighting COVID-19 exhausts T cells. Nat. Rev. Immunol. 20, 277.

Murira, A., and Lamarre, A. (2016). Type-I Interferon Responses: From Friend to Foe in the Battle against Chronic Viral Infection. Front. Immunol. 7, 609.

Ohno, Y., Kitamura, H., Takahashi, N., Ohtake, J., Kaneumi, S., Sumida, K., Homma, S., Kawamura, H., Minagawa, N., Shibasaki, S., et al. (2016). IL-6 down-regulates HLA class II expression and IL-12 production of human dendritic cells to impair activation of antigen-specific CD4+ T cells. Cancer Immunol. Immunother. 65, 193–204.

Parisi, G., Saco, J.D., Salazar, F.B., Tsoi, J., Krystofinski, P., Puig-Saus, C., Zhang, R., Zhou, J., Cheung-Lau, G.C., Garcia, A.J., et al. (2020). Persistence of adoptively transferred T cells with a kinetically engineered IL-2 receptor agonist. Nat. Commun. 11, 660.

Perry, A.K., Chen, G., Zheng, D., Tang, H., and Cheng, G. (2005). The host type I interferon response to viral and bacterial infections. Cell Res. 15, 407–422.

Remacle, F., Kravchenko-Balasha, N., Levitzki, A., and Levine, R.D. (2010). Information-theoretic analysis of phenotype changes in early stages of carcinogenesis. Proc. Natl. Acad. Sci. 107, 10324–10329.

Reyes-Sandoval, A., Berthoud, T., Alder, N., Siani, L., Gilbert, S.C., Nicosia, A., Colloca, S., Cortese, R., and Hill, A.V.S. (2010). Prime-boost immunization with adenoviral and modified vaccinia virus Ankara vectors enhances the durability and polyfunctionality of protective malaria CD8+ T-cell responses. Infect. Immun. 78, 145–153.

Ruan, Q., Yang, K., Wang, W., Jiang, L., and Song, J. (2020). Clinical predictors of mortality due to COVID-19 based on an analysis of data of 150 patients from Wuhan, China. Intensive Care Med. 46, 846–848.

Saeidi, A., Zandi, K., Cheok, Y.Y., Saeidi, H., Wong, W.F., Lee, C.Y.Q., Cheong, H.C., Yong, Y.K., Larsson, M., and Shankar, E.M. (2018). T-Cell Exhaustion in Chronic Infections: Reversing the State of Exhaustion and Reinvigorating Optimal Protective Immune Responses. Front. Immunol. 9, 2569.

Van de Sande, B., Flerin, C., Davie, K., De Waegeneer, M., Hulselmans, G., Aibar, S., Seurinck, R., Saelens, W., Cannoodt, R., Rouchon, Q., et al. (2020). A scalable SCENIC workflow for single-cell gene regulatory network analysis. Nat. Protoc. 15, 2247–2276.

Schmidl, C., Renner, K., Peter, K., Eder, R., Lassmann, T., Balwierz, P.J., Itoh, M., Nagao-Sato, S., Kawaji, H., Carninci, P., et al. (2014). Transcription and enhancer profiling in human monocyte subsets. Blood 123, e90–9.

Somers, E.C., Eschenauer, G.A., Troost, J.P., Golob, J.L., Gandhi, T.N., Wang, L., Zhou, N., Petty, L.A., Baang, J.H., Dillman, N.O., et al. (2020). Tocilizumab for treatment of mechanically ventilated patients with COVID-19. MedRxiv.

Stetson, D.B., and Medzhitov, R. (2006). Type I Interferons in Host Defense. Immunity 25, 373–381.

Sturm, G., Szabo, T., Fotakis, G., Haider, M., Rieder, D., Trajanoski, Z., and Finotello, F. (2020). Scirpy: A Scanpy extension for analyzing single-cell T-cell receptor sequencing data. BioRxiv 2020.04.10.035865.

Su, Y., Shi, Q., and Wei, W. (2017). Single cell proteomics in biomedicine: High-dimensional data acquisition, visualization, and analysis. Proteomics 17.

Su, Y., Bintz, M., Yang, Y., Robert, L., Ng, A.H.C., Liu, V., Ribas, A., Heath, J.R., and Wei, W. (2019). Phenotypic heterogeneity and evolution of melanoma cells associated with targeted therapy resistance. PLOS Comput. Biol. 15, 1–22.

Su, Y., Ko, M.E., Cheng, H., Zhu, R., Xue, M., Wang, J., Lee, J.W., Frankiw, L., Xu, A., Wong, S., et al. (2020). Multi-omic single-cell snapshots reveal multiple independent trajectories to drug tolerance in a melanoma cell line. Nat. Commun. 11, 2345.

Takeuchi, A., and Saito, T. (2017). CD4 CTL, a Cytotoxic Subset of CD4+ T Cells, Their Differentiation and Function. Front. Immunol. 8, 194.

Tan, L., Wang, Q., Zhang, D., Ding, J., Huang, Q., Tang, Y.-Q., Wang, Q., and Miao, H. (2020). Lymphopenia predicts disease severity of COVID-19: a descriptive and predictive study. Signal Transduct. Target. Ther. 5, 33.

van Ton, A.M., Kox, M., Abdo, W.F., and Pickkers, P. (2018). Precision Immunotherapy for Sepsis. Front. Immunol. 9, 1926.

Vardhana, S.A., and Wolchok, J.D. (2020). The many faces of the anti-COVID immune response. J. Exp. Med. 217.

Wang, Y., Liu, H., McKenzie, G., Witting, P.K., Stasch, J.-P., Hahn, M., Changsirivathanathamrong, D., Wu, B.J., Ball, H.J., Thomas, S.R., et al. (2010). Kynurenine is an endothelium-derived relaxing factor produced during inflammation. Nat. Med. 16, 279–285.

Wang, Z., Yang, B., Li, Q., Wen, L., and Zhang, R. (2020). Clinical Features of 69 Cases With Coronavirus Disease 2019 in Wuhan, China. Clin. Infect. Dis.

WHO (2020). COVID-19 Therapeutic Trial Synopsis, World Health Organization.

Wilk, A.J., Rustagi, A., Zhao, N.Q., Roque, J., Martínez-Colón, G.J., McKechnie, J.L., Ivison, G.T., Ranganath, T., Vergara, R., Hollis, T., et al. (2020). A single-cell atlas of the peripheral immune response in patients with severe COVID-19. Nat. Med.

Wittmann, B.M., Stirdivant, S.M., Mitchell, M.W., Wulff, J.E., McDunn, J.E., Li, Z., Dennis-Barrie, A., Neri, B.P., Milburn, M. V, Lotan, Y., et al. (2014). Bladder Cancer Biomarker Discovery Using Global Metabolomic Profiling of Urine. PLoS One 9, e115870.

Wolf, F.A., Angerer, P., and Theis, F.J. (2018). SCANPY: large-scale single-cell gene expression data analysis. Genome Biol. 19, 15.

Yan, L., Zhang, H.-T., Goncalves, J., Xiao, Y., Wang, M., Guo, Y., Sun, C., Tang, X., Jing, L., Zhang, M., et al. (2020). An interpretable mortality prediction model for COVID-19 patients. Nat. Mach. Intell. 2, 283–288.

Yang, Y., Shen, C., Li, J., Yuan, J., Yang, M., Wang, F., Li, G., Li, Y., Xing, L., Peng, L., et al. (2020). Exuberant elevation of IP-10, MCP-3 and IL-1ra during SARS-CoV-2 infection is associated with disease severity and fatal outcome. MedRxiv.

Zadran, S., Arumugam, R., Herschman, H., Phelps, M.E., and Levine, R.D. (2014). Surprisal analysis characterizes the free energy time course of cancer cells undergoing epithelial-to-mesenchymal transition. Proc. Natl. Acad. Sci. 111, 13235–13240.

Zheng, G.X.Y., Terry, J.M., Belgrader, P., Ryvkin, P., Bent, Z.W., Wilson, R., Ziraldo, S.B., Wheeler, T.D., McDermott, G.P., Zhu, J., et al. (2017). Massively parallel digital transcriptional profiling of single cells. Nat. Commun. 8, 14049.

Zheng, M., Gao, Y., Wang, G., Song, G., Liu, S., Sun, D., Xu, Y., and Tian, Z. (2020). Functional exhaustion of antiviral lymphocytes in COVID-19 patients. Cell. Mol. Immunol. 17, 533–535.

Zhou, J., Zhou, S., and Zeng, S. (2013). Experimental diabetes treated with trigonelline: effect on β cell and pancreatic oxidative parameters. Fundam. Clin. Pharmacol. 27, 279–287.

Zhou, J., Kaiser, A., Ng, C., Karcher, R., McConnell, T., Paczkowski, P., Fernandez, C., Zhang, M., Mackay, S., and Tsuji, M. (2017). CD8+ T-cell mediated anti-malaria protection induced by malaria vaccines; assessment of hepatic CD8+ T cells by SCBC assay. Hum. Vaccin. Immunother. 13, 1625–1629.

Zhou, Y., Fu, B., Zheng, X., Wang, D., Zhao, C., qi, Y., Sun, R., Tian, Z., Xu, X., and Wei, H. (2020). Aberrant pathogenic GM-CSF+ T cells and inflammatory CD14+CD16+ monocytes in severe pulmonary syndrome patients of a new coronavirus. BioRxiv.

